# Spatio-Temporal Coordination of Transcription Preinitiation Complex Assembly in Live Cells

**DOI:** 10.1101/2020.12.30.424853

**Authors:** Vu Q. Nguyen, Anand Ranjan, Sheng Liu, Xiaona Tang, Yick Hin Ling, Jan Wisniewski, Gaku Mizuguchi, Kai Yu Li, Vivian Jou, Qinsi Zheng, Luke D. Lavis, Timothée Lionnet, Carl Wu

## Abstract

Transcription initiation by RNA polymerase II (Pol II) requires <u>p</u>reinitiation <u>c</u>omplex (PIC) assembly at gene promoters. In the dynamic nucleus where thousands of promoters are broadly distributed in chromatin, it is unclear how ten individual components converge on any target to establish the PIC. Here, we use live-cell, single-molecule tracking in *S. cerevisiae* to document subdiffusive, constrained exploration of the nucleoplasm by PIC components and Mediator’s key functions in guiding this process. On chromatin, TBP, Mediator, and Pol II instruct assembly of a short-lived PIC, which occurs infrequently but efficiently at an average promoter where initiation-coupled disassembly may occur within a few seconds. Moreover, PIC exclusion by nucleosome encroachment underscores regulated promoter accessibility by chromatin remodeling. Thus, coordinated nuclear exploration and recruitment to accessible targets underlies dynamic PIC establishment in yeast. Collectively, our study provides a global spatio-temporal model for transcription initiation in live cells.

## INTRODUCTION

RNA polymerase II (Pol II) transcribes the vast majority of eukaryotic genes. To initiate this process, Pol II associates with general transcription factors (GTFs) TFIID, <u>T</u>ATA-<u>b</u>inding <u>p</u>rotein TBP, TFIIA, TFIIB, TFIIE, TFIIF, TFIIH, TFIIK and the coactivator Mediator to establish a <u>p</u>re-initiation <u>c</u>omplex (PIC) at gene promoters (Cramer, 2019; Robinson et al., 2016; Schilbach et al., 2017). Conserved DNA elements are directly recognized by TFIID and TBP in the presence of TFIIA (Patel et al., 2018; Zhang et al., 2016), while Pol II and remaining GTFs are generally tasked with enzymatic activities for promoter melting and scanning for the transcription start site (TSS) (Cramer, 2019). The Mediator may dynamically transit between upstream activating sequences (UASs) and promoters in yeast (Jeronimo et al., 2016; Knoll et al., 2018; Petrenko et al., 2016; Wong et al., 2014) and form puncta or phase-separated condensates with sequence-specific transcription factors (TFs) at super-enhancers in ESCs (Boija et al., 2018; Shrinivas et al., 2019), but its dynamics and specific roles in transcription initiation are not fully understood (Khattabi et al., 2019; Knoll et al., 2018; Petrenko et al., 2017). While PIC establishment has been extensively studied *in vitro*, the mechanism and kinetics underlying this process *in vivo* remain fundamental questions in gene regulation.

Individual PIC components searching for promoters must contend with multiple levels of genome organization in the nucleus. PIC components tend to bind nucleosome-depleted regions (NDRs) adjoining the downstream “+1” nucleosome genome-wide (Lauberth et al., 2013; Rhee and Pugh, 2012), and this association is sensitive to characteristic histone modifications (Joo et al., 2017; Kubik et al., 2018; Lauberth et al., 2013; Vermeulen et al., 2007). Thus, proximal nucleosomes contribute to the recognition of native promoters, many of which lack core promoter motifs (Haberle and Stark, 2018). The PIC must also negotiate promoter access with multiple chromatin regulators recruited to the NDRs to mobilize and modify local nucleosomes. Beyond gene promoters, non-specific interactions with the vast excess of bulk chromatin can affect target-search efficiency of PIC components (Normanno et al., 2012). Furthermore, the abundances and distributions of individual components also influence PIC formation. Despite stoichiometric composition within the complex, PIC components are present at different concentrations (Borggrefe et al., 2001; Kimura et al., 1999) and at least two—Mediator and Pol II—have been shown to traffic in transient and stable clusters in mammalian cells (Cho et al., 2018; Cisse et al., 2013; Li et al., 2019). Understanding PIC establishment *in vivo* demands quantitative knowledge of how individual components explore the nucleoplasm and chromatin to find promoter targets.

Single-molecule tracking (SMT) is a powerful approach to characterize the dynamic behaviors of factors in live cells at high spatiotemporal resolution (Mazza et al., 2012). SMT enables visualization of nuclear factors diffusing in space, transiently sampling chromatin and stably associating with specific targets, informing various modes of target search (Hansen et al., 2020; Izeddin et al., 2014). This dynamic view complements steady-state snapshots acquired by genomics and static structural approaches. However, despite increasing SMT implementation, systematic investigations of factors that assemble a functional macromolecular complex on chromatin are limited, precluding our understanding of this essential process in genome metabolism.

Here, we apply live-cell SMT in budding yeast to capture the dynamics of all ten PIC components in the nucleoplasm and on chromatin. By combining SMT with conditional depletion experiments, we examine different modes of spatial exploration in the nucleoplasm and hierarchical recruitment of PIC components to chromatin. Furthermore, we determine target residence times and extrapolate other parameters relevant to transcription initiation kinetics in live cells. Our findings suggest a model in which spatio-temporal coordination among key factors enables convergence of PIC components for transcription initiation on a timescale of several seconds.

## RESULTS

### Global dynamics of PIC components in live cells

To investigate PIC establishment at gene promoters *in vivo*, we first characterized the global diffusive behaviors and chromatin association of all ten PIC components. We created individual strains, each expressing as the sole source a terminal fusion of HaloTag (Los et al., 2008) on Rpb1 (Pol II), Med14 (Mediator), Taf1 (TFIID), TBP, Toa1 (TFIIA), Sua7 (TFIIB), Tfg1 (TFIIF), Tfa1 (TFIIE), Ssl2 (TFIIH), or Kin28 (TFIIK) (Figure 1A). Wildtype-level growth of these strains confirmed functionality of the tagged, essential proteins (Figures S1A-C). We performed all imaging experiments on cells undergoing logarithmic growth in rich glucose media; accordingly, our observations relate on average to the predominant transcription of major housekeeping genes (Pelechano et al., 2010).

**Figure 1.**
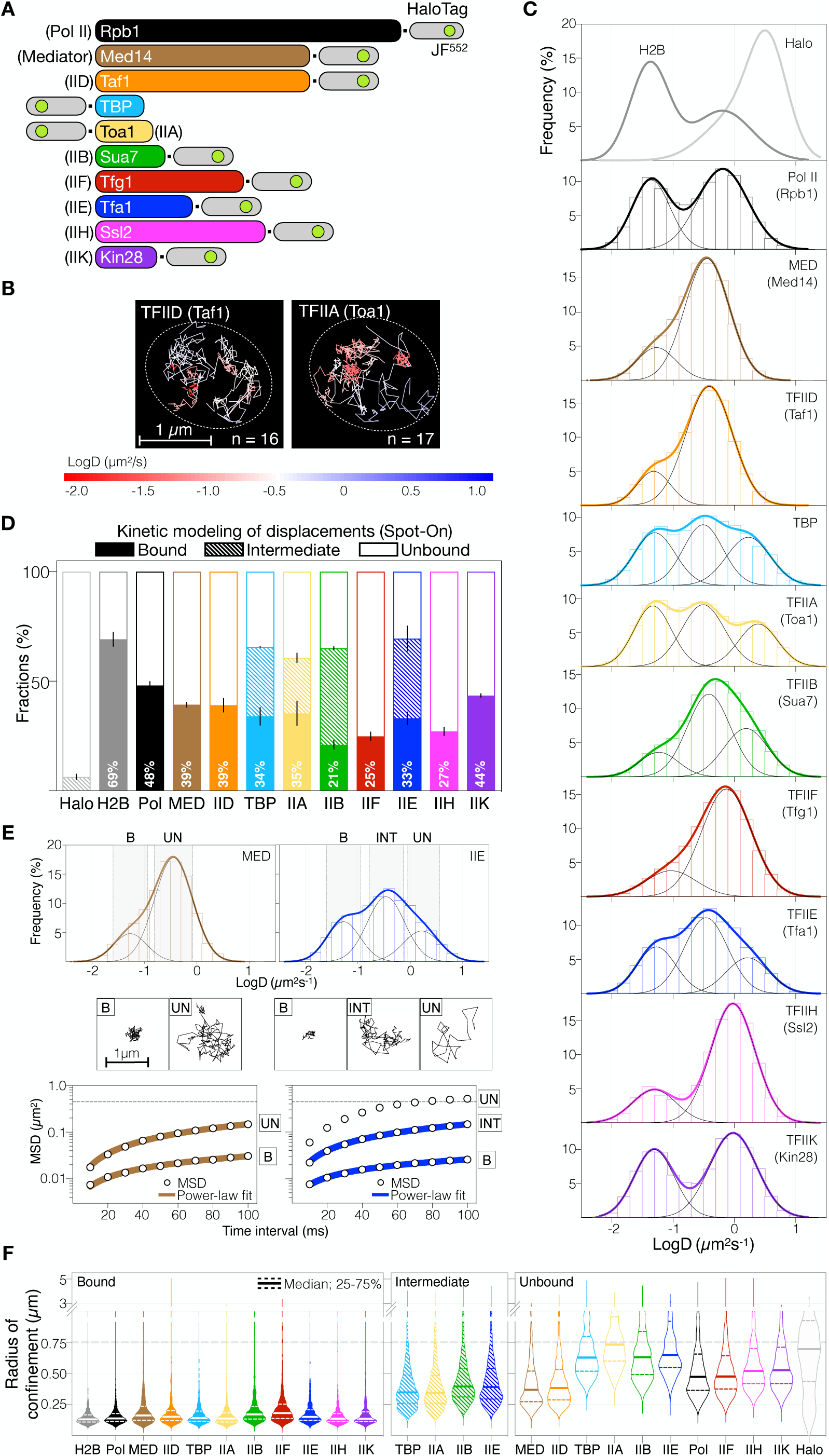
Global dynamics of individual PIC components. (A) Schematics of N- or C-terminal HaloTag fusions to scale in individual yeast strains. JF^552^, fluorescent ligand Janelia Fluor 552. (B) Representative overlays of single-molecule TFIID and TFIIA trajectories, colored according to calculated diffusion coefficients. Nuclei are demarcated (ovals). N, number of trajectories presented. (C) Diffusion coefficient (D, log_10_) histograms (bars) and multi-Gaussian fits (thick curves) of H2B and nuclear Halotag (top, only fits shown) and PIC components, where resolved populations are shown (thin black curves). Histograms contain data from three biological replicates. (D) Relative frequencies of dynamic behaviors acquired from kinetic modeling (Figure S1E and Table S1), with % chromatin binding (F_B_) indicated. Results are means ± SD from three biological replicates. (E) Top: fitted MED and TFIIE logD histograms as in (B). Trajectories exhibiting logD values within mean ± SD of respective population (shaded bars) were selected. B, chromatin-bound. UN, unbound. INT, intermediate. Middle: representative MED and IIE single-molecule trajectories. Bottom: average mean-squared displacement (MSD) of each population with corresponding power-law fit. Fit results are plotted in Figure S2C. (F) Violin plots of radii of confinement (Rc, µm) calculated for single trajectories of bound, intermediate and unbound populations (Figure S2 and see Methods). Rc = 0.75 µm (horizontal dashed line) represents the yeast nuclear radius as indicated by the median Rc of HaloTag.

We labeled HaloTag fusions with Janelia Fluor 552 (JF^552^) (Zheng et al., 2019), imaged single molecules at 10 ms resolution and tracked their positions with ∼30 nm x-y precision to establish 2D projections of trajectories through nuclear space (Figure 1B, see Methods). We implemented this “fast-tracking” regime to acquire 10,000-50,000 trajectories for each PIC component as well as histone H2B and nuclear HaloTag, which served as live-cell standards for chromatin-bound and unbound behaviors, respectively. We calculated the mean-squared displacement (MSD) for each trajectory to extract the molecule’s apparent diffusion coefficient (D) (see Methods). The logD distributions showed values within the technical dynamic range demonstrated by histone H2B and free HaloTag (Figures 1C and S1D). Gaussian fitting resolved two dynamically distinct populations for Pol II, Mediator, TFIID, IIF, IIH and IIK, and unexpectedly, three for TBP, TFIIA, IIB and IIE (Figure 1C). Accordingly, we performed two- or three-state kinetic modeling of single displacements (Hansen et al., 2018) to achieve more robust quantitation of the average D (av. D) and fraction of molecules (F) comprising each population (Figures 1D, S1E and Table S1, see Methods).

As anticipated, the majority of H2B (F∼70%) exhibited low mobility (av. D=0.01 µm^2^/s), taken as the average diffusivity of yeast chromatin (Figures 1C and 1D). However, the lowest-mobility populations of PIC components, assigned as chromatin-bound and subsequently validated using binding mutants (below), displayed several-fold faster diffusion (av. D=0.03-0.06 µm^2^/s), suggesting that genomic sites bound by PIC components are generally more dynamic than bulk chromatin. Importantly, the relatively minor fractions of bound Mediator and all GTFs (F_B_=21%-44%) indicate that most molecules underwent diffusive processes in the nucleoplasm. Note that F_B_ values represent all binding at promoter targets and non-specific sites throughout chromatin. F_B_ for Pol II (48%) also includes elongating and terminating molecules that engaged downstream of promoters, while that for TBP (34%) includes binding at Pol I and Pol III-transcribed genes.

At the other extreme, diffusivities of the most mobile populations (av. D=0.6-3.3 µm^2^/s), assigned as unbound molecules, correlated negatively with the molecular weights (MW) of individual PIC components (Figure S1F), as expected of molecules diffusing in the same medium. Notably, unbound H2B diffused more slowly (av. D=1.6 µm^2^/s) than expected (>2 µm^2^/s) for a species of similar MW to H2A-Halo-H2B (∼55 kDa), consistent with its biochemical association with various histone chaperones in the nucleoplasm (Hammond et al., 2017). The low mobilities of Med14 and Taf1 (av. D=0.6 µm^2^/s) suggest that they traverse the nucleoplasm in large Mediator and TFIID complexes of up to 1 MDa MW. Intriguingly, the small GTFs TBP, TFIIA, IIB and IIE, whose unbound populations were the most mobile among PIC components (av. D > 2 µm^2^/s) (Figure S1F), also displayed a well-resolved, less mobile third population with diffusivities (av. D=0.4-0.6 µm^2^/s) resembling those of unbound Mediator and TFIID (Figures 1C, 1D and Table S1). This “intermediate” population occurred at substantial fractions (F_I_=25%-44%), and was absent or unresolved for the other PIC components (Pol II, TFIIF, IIH, and IIK), indicating that this is not general behavior within the yeast nucleus. Considered together, our results indicate that highly dynamic PIC components infrequently interact with chromatin and more often undergo diffusive processes to explore the nucleoplasm.

### Subdiffusive exploration of nuclear space by PIC components

To acquire a spatial perspective on the three dynamic behaviors of PIC components, we sub-classified trajectories into bound, intermediate and unbound populations according to Gaussian fitting of the logD histograms (Figures 1E and S2A). The ensemble MSDs computed for bound populations of PIC components and H2B were well fit by a power law model (MSD ∝ t^α^) with anomalous coefficient α=0.5-0.6 (Figures S2B and S2C) characteristic of subdiffusion (α <1) (Woringer and Darzacq, 2018; Woringer et al., 2020). These results are quantitatively consistent with the reported mobility of a chromosomal *tetO* array in yeast (Miné-Hattab et al., 2017). To estimate the spatial constraint imposed by chromatin, we fit the MSDs of individual trajectories with a circular-confined diffusion model to extract the radius of confinement (Rc) (Wieser and Schütz, 2008) as recently demonstrated for mammalian H2B and chromatin-binding factors (Lerner et al., 2020). As expected, the Rc distributions are similar among bound populations of PIC components and H2B, with median Rc∼0.13 µm representing the spatial confinement of chromatin in yeast (Figure 1F).

Remarkably, intermediate populations of TBP, TFIIA, IIB and IIE produced ensemble MSDs that also fit the power-law function with α=0.8 indicative of a subdiffusive behavior distinct from that of chromatin (Figures S2B and S2C). Rc calculations suggest that molecules displaying this behavior tend to explore sub-nuclear areas with similar apparent median Rc∼0.4 µm (the radius of the yeast nucleus is ∼0.75 µm) (Figure 1F). By contrast, ensemble MSDs from the unbound populations of these four GTFs are characteristic of free diffusion, displaying plateaus consistent with confinement to the average dimensions of the yeast nucleus (Figure S2B) (Dion and Gasser, 2013). Thus, the three dynamic populations resolved for TBP, TFIIA, IIB and IIE correspond to chromatin-bound, subdiffusive and free diffusive states.

Unbound Mediator and TFIID also displayed sub-diffusive dynamics with α=0.9 (Figures S2B and S2C) and median Rc∼0.4 µm, notably similar to the intermediate populations of TBP, TFIIA, IIB and IIE (Figure 1F). To further investigate the manner by which these six PIC components explore apparently limited space, we analyzed the spatial angles formed by consecutive displacements of single molecules (Figure S2D) (Hansen et al., 2020). Nuclear trajectories of unbound Mediator and TFIID, as well as intermediate populations of TBP, TFIIA, IIB and IIE exhibited almost two-fold enrichment (1.3-fold for TBP) of large angles (180°±30°) relative to small angles (0°±30°) (Figure S2E). This directional bias would manifest a back-and-forth trajectory consistent with confined diffusion whereby molecules oversample a local environment, such as a transient “trapping zone” proposed for mammalian CTCF (Hansen et al., 2020; Izeddin et al., 2014). Importantly, unbound TBP, TFIIA, IIB and IIE displayed no such anisotropic angle distributions (Figures S2D and S2E), indicating their unconstrained mobility within the yeast nucleus.

In addition, unbound Pol II, TFIIF, IIH and IIK also exhibited subdiffusion with α=0.8 (Figures S2B and S2C) and median Rc∼0.5 µm (Figure 1F). Notably, the indistinguishable diffusive behaviors of unbound Pol II and TFIIF (Figures 1F, S1F and S2C) suggest that they associate in the nucleoplasm, consistent with biochemical properties of the two factors in yeast nuclear extracts (Rani et al., 2004). Similarly, the diffusive behaviors of TFIIH and IIK are also consistent with their biochemical interactions (Keogh et al., 2002). Taken together, our analysis reveals that all PIC components undergo subdiffusive processes to explore the yeast nucleus, with a tendency to sample 0.4-0.5 µm-radial areas.

### TBP, Mediator and Pol II instruct hierarchical PIC assembly

Purified PIC components biochemically engage promoter DNA in a stepwise fashion to assemble a PIC *in vitro*, but it is unclear how PIC components diffusing in the nucleoplasm are recruited to chromatin in live cells. We sought to deconstruct this process by removing key factors to determine how they influence chromatin binding by the remaining PIC components. To explore the role of TBP, the key factor that nucleates genome-wide assembly of the PIC, we used the Anchor-Away (AA) technique to conditionally deplete it from the nuclei of growing cells (Haruki et al., 2008). Efficient depletion was achieved after one-hour treatment with rapamycin, which coupled ribosome processing with eviction of TBP tethered to the ribosomal protein Rpl13A *via* FRB and FKBP12 tags, respectively (Figure S3A). Other PIC components remained nuclear with little to no observable co-depletion, allowing fast-tracking experiments to assess overall chromatin binding in the singular absence of TBP. We found that AA of TBP reduced chromatin-binding by Pol II, TFIIA, IIH, and IIK to minute, unquantifiable levels (Figures 2A, 2B, S3B, S3C and S3D). Notably, along with chromatin binding, subdiffusion by TFIIA was nearly abolished (Figure 2A), thus linking this mode of spatial exploration to the promoter-recognition process by TFIID, TBP and TFIIA (Kraemer et al., 2001; Patel et al., 2018). By contrast, we found increased binding by Mediator (+20%) and especially TFIID (+70%), consistent with genome-wide binding at UASs and promoters, respectively, under similar TBP depletion conditions (Knoll et al., 2018, 2020). The dramatic response by TFIID also implicates an inhibitory effect of TBP on TFIID binding at promoters, as recently shown *in vitro* (Le et al., 2019).

**Figure 2.**
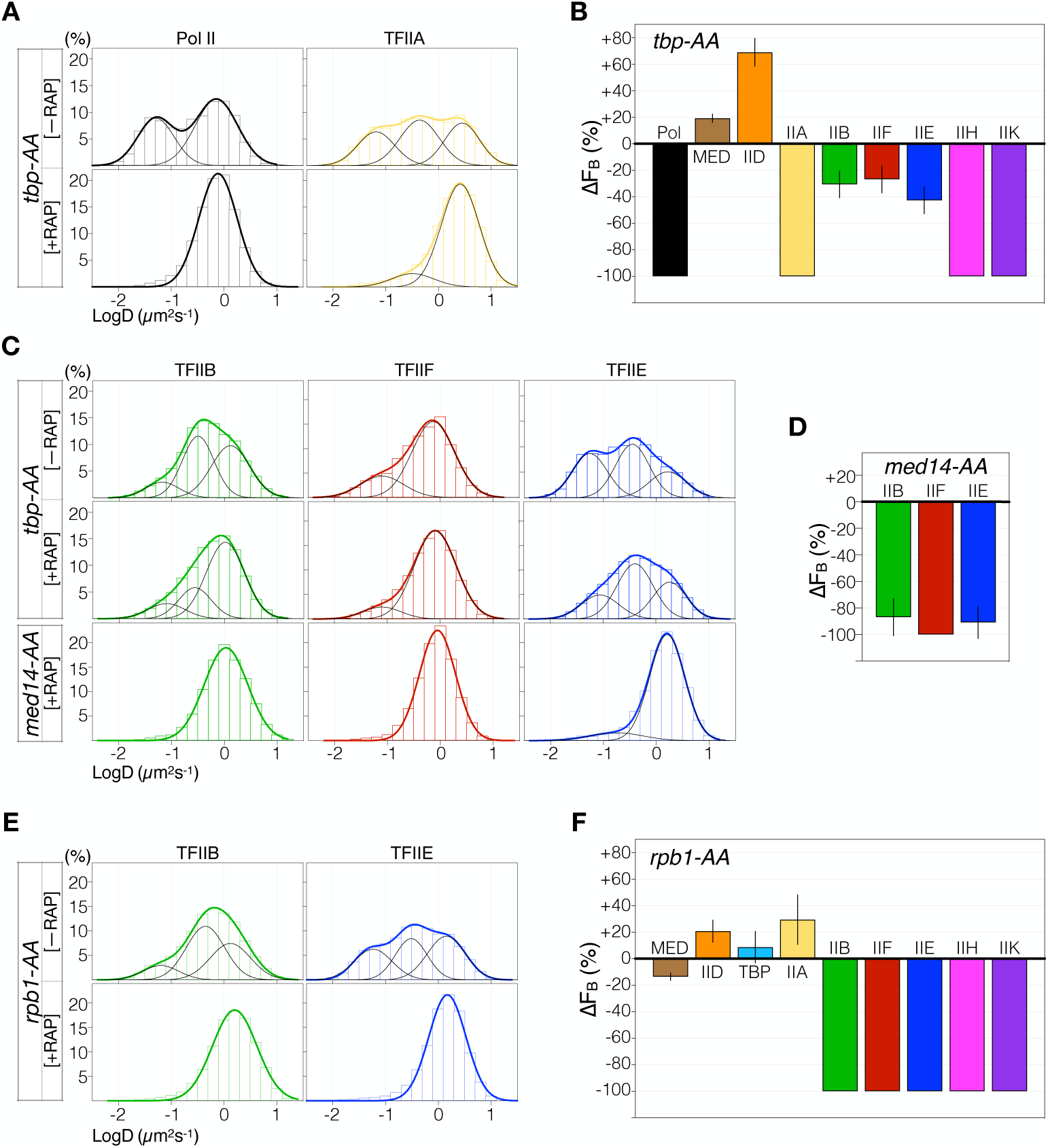
TBP, Mediator and Pol II instruct hierarchical PIC assembly. (A) LogD histograms of Pol II and TFIIA before (-RAP, top) and after (+RAP, bottom) TBP depletion by rapamycin-dependent Anchor Away (*TBP AA*). Gaussian fits (thick curves) and resolved populations (thin curves) are shown. Histograms contain data from one (-RAP) and three (+RAP) biological replicates. (B) Changes in F_B_ (ΔF_B_) relative to wildtype after TBP depletion, calculated using mean F_B_ ± SD (Figure S3D and Table S2), from three biological replicates of each condition (see Methods), and errors were propagated. [One *TBP AA* sample without rapamycin was imaged to confirm wildtype-level F_B_ (Figures S3C, S3D and Table S2)]. (C) LogD histograms of TFIIB, IIF and IIE before (-RAP, top row) and after (+RAP) TBP depletion (*TBP AA*, middle row) or Med14 depletion (*MED14 AA*, bottom row). Data are shown as in (A). (D) ΔF_B_ relative to wildtype after Med14 depletion, computed and shown as in (B). (E) LogD histograms of TFIIB and IIE in *RPB1 AA* cells before (-RAP, top) and after (+RAP, bottom) Pol II depletion, shown as in (A). (F) ΔF_B_ relative to wildtype after Rpb1 depletion, computed and shown as in (B).

Unexpectedly, after TBP depletion, we observed lower binding by TFIIB, IIE and IIF (27% to 43% decreases) (Figures 2B and 2C). We next sought to determine whether Mediator was responsible for chromatin binding by these three GTFs, as it has been shown to recruit TFIIB to promoter DNA in HeLa extracts by *in vitro* biochemistry (Baek et al., 2006). Remarkably, AA depletion of Med14, a critical structural subunit of Mediator, resulted in full or near-complete abolishment of chromatin binding by TFIIB (84% decrease), TFIIE (92% decrease) and TFIIF (100% decrease) (Figures 2C and 2D), indicating Mediator’s dominant role in recruiting TFIIB, IIE and IIF. Double depletion of TBP and Med14 elicited identical responses (Figures S3E and S3F), indicating that chromatin binding by TFIIB, IIE and IIF observed after TBP depletion (Figure 2C) was facilitated by Mediator. The partial TBP requirement observed for this process (Figures 2B and 2C) may reflect its role in Mediator association at a small subset of promoters (Knoll et al., 2018).

TFIIB, IIE and IIF physically interface with the PIC *via* Pol II (Robinson et al., 2016; Schilbach et al., 2017). In light of Mediator’s critical role in recruiting these three GTFs, we also explored Pol II’s involvement by AA depletion of its catalytic subunit Rpb1. As expected, Rpb1 co-depleted with another Pol II subunit Rpb9 and no other PIC component (Figure S3G). Under Rpb1 AA, we observed a modest effect on chromatin binding by Mediator, while that by TFIIB, IIE, IIF, IIH and IIK—all enzymatic components of the PIC—was essentially abolished (Figures 2E, 2F, S3H and S3I). Thus, both Mediator and Pol II are required for the recruitment of TFIIB, IIE and IIF. This dual requirement applies to both subdiffusion and chromatin binding of TFIIB and IIE, as these two behaviors were nearly ablated upon either Med14 (Figure 2C) or Rpb1 depletion (Figure 2E). Our observations suggest that subdiffusion in the nucleoplasm provides a conducive environment for recruitment of TFIIB and IIE to chromatin, and that both processes are coordinated by Mediator and Pol II. Importantly, Rpb1 depletion elicited only modest changes in binding by TFIID, TBP and TFIIA, similar to Mediator (Figures 2F, S3H and S3I) and consistent with genome-wide ChIP studies (Joo et al., 2017; Knoll et al., 2018). These results support a model in which Mediator and TFIID-TBP-TFIIA together organize a platform at promoters for stepwise recruitment of Pol II and the remaining GTFs, as shown *in vitro* (Johnson and Carey, 2003).

Our AA experiments affirm TBP’s critical role in PIC assembly and reveal a previously uncharacterized collaboration between Mediator and Pol II to coordinate recruitment of the enzymatic components of the PIC involving subdiffusive exploration by TFIIB and IIE. Furthermore, disruption of this dynamic hierarchy by AA depletions abolished chromatin binding for seven PIC components (Pol II, TFIIA, TFIIB, IIE, IIF, IIH and IIK), suggesting a remarkable absence of detectable ‘non-specific’ chromatin interactions beyond promoter regions. Importantly, based on the strict dependencies demonstrated here, chromatin binding detected for the enzymatic GTFs also indicates presence of TBP, Mediator and Pol II in the bound entity taken as the PIC. This interpretation circumvents technical limitations on multi-color SMT imaging for direct PIC observation at any one location in live cells.

### Short-lived PIC resides for only a few seconds on promoter chromatin

The dynamic nature of the PIC has been inferred from genome-wide ChIP studies (Wong et al., 2014), and biochemical experiments have demonstrated that most PIC components disengage from the complex prior to promoter escape *in vitro* (Fujiwara et al., 2019). However, their *in* vivo residence times on chromatin, which should inform the lifetime of the PIC and provide key metrics for transcription initiation kinetics, are unknown. To determine these parameters, we first performed “slow tracking” to selectively visualize and determine the dwell times of binding events. This imaging regime features long exposure time (250 ms) to blur out diffusing molecules and low laser power to limit fluorophore photobleaching (Chen et al., 2014). Individual survival probability curves (1-CDF), computed from the observed dwell times of several thousand binding events, showed that on average, all PIC components, except Pol II and TBP, were substantially shorter-lived on chromatin than H2B (Figure 3A), which indicated the temporal limit imposed by photobleaching and chromatin motions (Hansen et al., 2017). To accommodate longer dwell times of Pol II and TBP, we sought to further reduce the photobleaching limit by time-lapse imaging, which incorporates alternating excitation laser “on” (250 ms) and “off” (250-500 ms) periods (Gebhardt et al., 2013). This approach allowed better separation of the survival curve of Pol II from the bulk histone H2B standard for long-term chromatin residence (Figure S4E). However, time-lapse did not allow distinction between TBP and H2B, indicating that the factor’s dwell time is beyond resolution by SMT.

**Figure 3.**
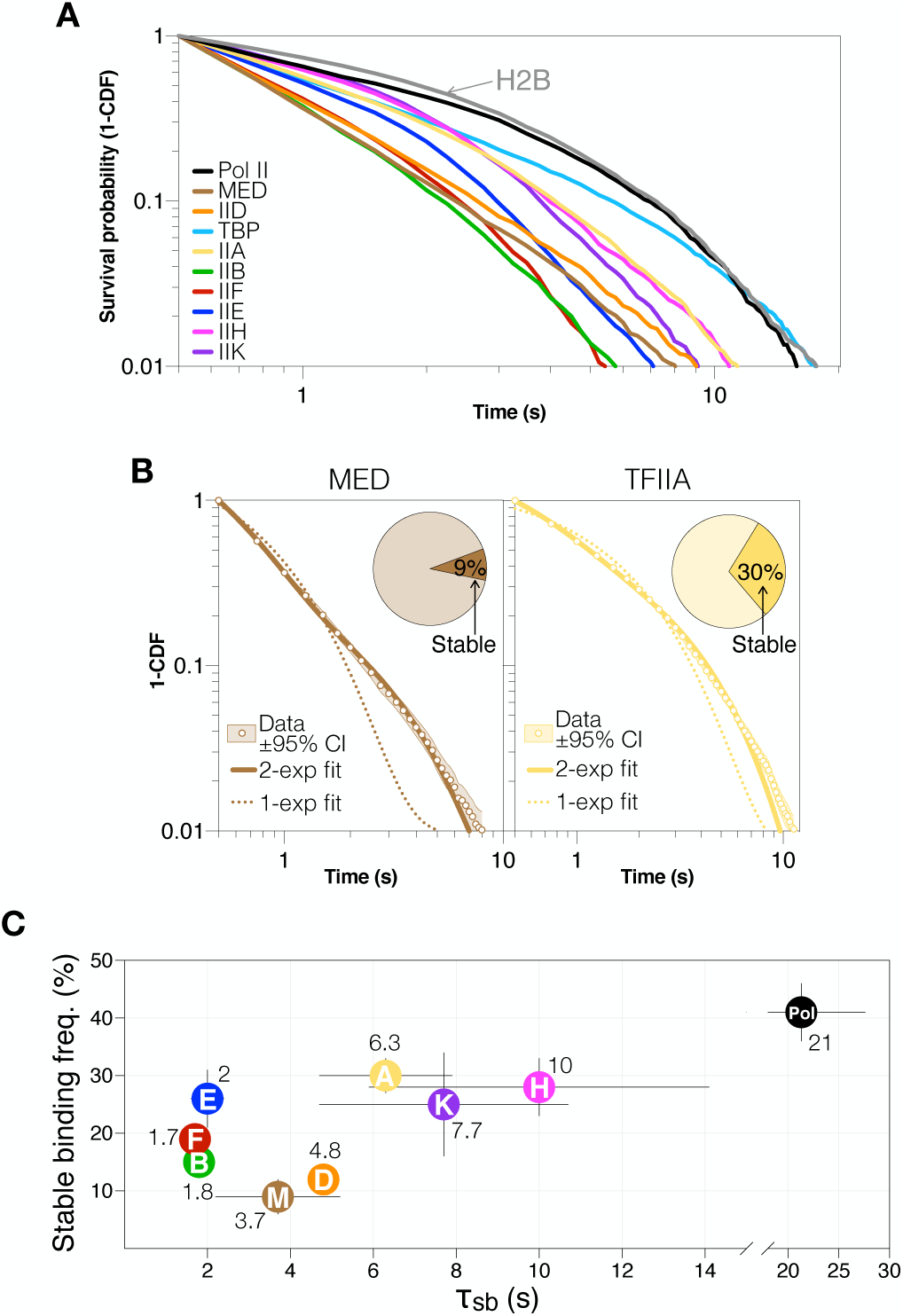
The assembled PIC is a short-lived entity in live cells. (A) Log-log survival-probability curves (1-CDF) from apparent dwell times of single-molecule chromatin-binding events. Curves contain data from three biological replicates. (B) 1-CDF data (white dots) of Mediator and TFIIA from (A), with ±95% confidence interval (CI) obtained by resampling (shaded area). Double-exponential fit (solid line) indicated fractions of molecules engaged in stable binding (f_sb_) and transient binding (pie chart, f_sb_ values shown) and respective apparent dissociation rates (k_sb_ and k_tb_, reported in Table S1). Single-exponential fit (dotted line) is shown for each dataset. (C) f_sb_ (%) and average residence times τ_sb_(s) of stable chromatin association, obtained by double-exponential fitting of the 1-CDF curves and H2B correction (Figure S5D, Table S1 and see Methods). τ_sb_ values are indicated. Results are means ± SD from three biological replicates. Values for Pol II were obtained by time-lapsed SMT imaging (Figure S5E).

Within the SMT resolvable range, each survival curve for PIC components minimally required a double-exponential fit corresponding to stable and transient binding characterized by average dissociation rates (k_sb_ and k_tb_, respectively) and fractions of all binding events (f_sb_ and f_tb_) (Figures 3B and S4D, Table S1). We used H2B decay kinetics to correct k values (Figure S4C, see Methods) (Hansen et al., 2017) and computed the average residence times (τ =1/k) for stable (τ_sb_) and transient (τ_tb_) binding (Figure 3C and Table S1). Both time-lapse regimes (250 ms and 500 ms “off”) produced similar of 20-23 s for Pol II (fsb ∼40%), τ_sb_ consistent with a value of 26 s derived by fluorescence recovery after photobleaching (FRAP) (Figures S4F and S4G). Among GTFs, TFIIA, IIH and IIK exhibited the longest τ_sb_, ranging between 6 s and 10 s (f_sb_=25-30%) (Figure 3C). Mediator and TFIID displayed τ_sb_ of 4-5 s (f_sb_∼10%, lowest among all components). The shortest-lived components TFIIB, IIE and IIF showed τ_sb_ ∼ 2 s (f_sb_=15%-25%), indicating that on average, a full PIC assemblage lasts only 2 seconds on chromatin in living cells.

Importantly, TBP depletion resulted in miniscule fractions of detectable binding events (1-2%) and even shorter residence times for Pol II (2 s), TFIIA (4s), IIH and IIK (3s) (Figures S4H and S4I), confirming the link between their normal τ_sb_ to PIC formation at promoters and Pol II transcription. For TFIIB, IIE, and IIF, TBP depletion had little effect on τ_sb_ but reduced f_sb_ by ∼50% (Figures S4H and S4I). Because we observed that recruitment of these three factors requires Pol II (Figure 2F), the miniscule Pol II fraction with a similar residence time (∼2 s) under TBP depletion may be an indication of that interaction with TFIIB, IIE, and IIF. Finally, consistent with fast-tracking results, TBP depletion had little effect on Mediator but substantially increased the τ_sb_ and f_sb_ for TFIID (Figures S4H and S4I). Together, our findings provide direct, quantitative metrics on the dynamic interactions of PIC components with promoter chromatin in the yeast nucleus.

### Kinetic coupling between promotor escape and PIC lifetime

To explore a connection between the short lifetime of the PIC and the transcription initiation process in live cells, we disrupted a key initiation event between PIC assembly and Pol II escape, involving serine-5 phosphorylation of the Rpb1 C-terminal domain (CTD) by the Kin28 kinase (TFIIK). Kin28 depletion or inactivation is associated with delayed Pol II promoter escape and global reduction of nascent RNA synthesis (Rodríguez-Molina et al., 2016; Wong et al., 2014). If transcription initiation were driving PIC turnover in live cells, its delay should result in longer dwell times for PIC components. Indeed, we found that Kin28 depletion (Figure S5A) resulted in at least 2-fold longer *τ*_sb_ for Mediator and all GTFs except

TFIIH (Figures 4A, 4B, S5B and S5C), thus linking their dissociation to the presence of Kin28 in wildtype cells. The shorter TFIIH *τ*_sb_ suggests that its stable PIC engagement involves an interaction interface provided by TFIIK, as demonstrated by cryo-EM structures (Schilbach et al., 2017). In addition, upon Kin28 depletion, Pol II exhibited a substantially shorter residence time of 10 s (compared to 23 s in wildtype cells) (Figures 4A and 4B). Complementary to ChIP and transcriptomics studies (Rodríguez-Molina et al., 2016; Wong et al., 2014), the 10 s residence time of Pol II represents an average of heterogeneous molecules stalled near promoters and elongating or prematurely terminated along gene bodies. To further validate these results, we performed fast tracking after Kin28 depletion (Figures 4C, S5D and S5E). As expected from their altered dissociation kinetics, we observed larger chromatin-bound fractions for Mediator, TFIID, TBP, TFIIA, IIB and IIE (20% to 80% increase), in contrast to Pol II (65% decrease) and TFIIH (30% decrease). Of interest, despite displaying longer *τ*_sb_, global binding by TFIIF was modestly decreased (by 20%) under Kin28 depletion, which may reflect a global reduction in elongation complexes where TFIIF, unlike other PIC components, has been shown to be present and functional (Schweikhard et al., 2014; Zawel et al., 1995).

**Figure 4.**
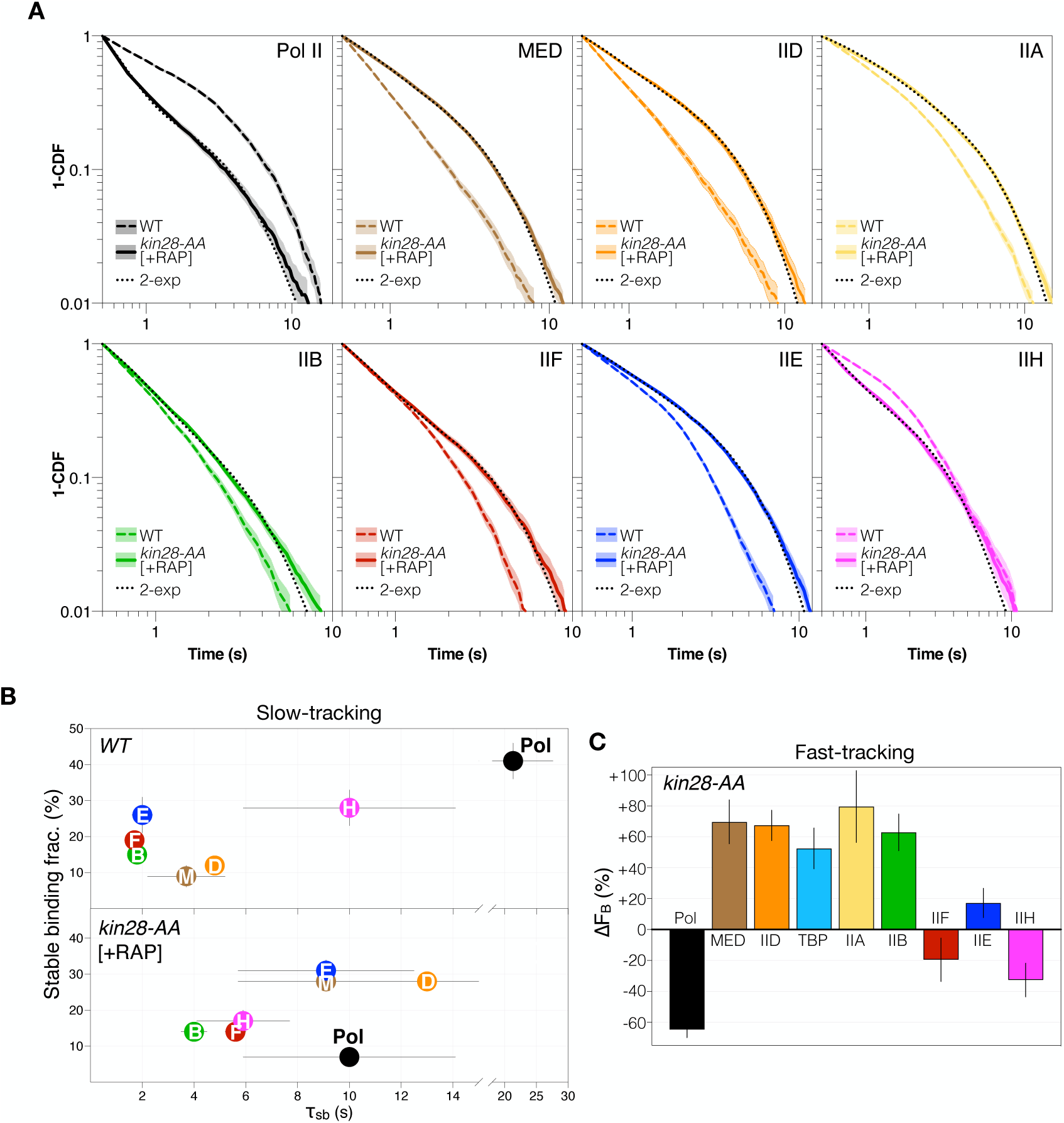
Delayed chromatin dissociation upon Kin28 depletion. (A) 1-CDF data and double-exponential fit obtained for PIC components after Kin28 depletion, shown as in Figure 3B. Wildtype data (WT, without fit) is shown for comparison. [One *KIN28 AA* sample without rapamycin was imaged to confirm wildtype-level dissociation kinetics (Figure S5D and Table S2).] Curves contain data from three biological replicates. (B) f_sb_ (%) and τ_sb_(s) of stable association by PIC components, obtained by slow tracking in wildtype cells (*WT*, top) and after Kin28 depletion (+RAP, bottom). [Values for TFIIA are not reported because its extended survival approached the H2B limit (Figure S5B).] All values are means ± SD from three biological replicates. Double-exponential fit parameters are reported in Table S2. (C) ΔF_B_ relative to wildtype after Kin28 depletion, obtained by fast tracking and shown as in Figure 2B. LogD histograms and kinetic modeling results are reported in Figures S5D, S5E and Table S2.

These results extend upon genome-wide ChIP findings for Mediator and TFIID under Kin28-AA conditions (Knoll et al., 2020; Wong et al., 2014) and further suggest the emergence of a temporally stalled PIC. Therefore, transcription initiation mediated by Kin28/TFIIK is a major determinant of rapid PIC turnover in live cells.

### Highly dynamic target search and low promoter occupancy by PIC components

We next sought to reconstruct the average trajectory of a molecule undergoing target search (Figure 5A). The duration of this process, or the τ_search_, is the temporal sum of alternating transient chromatin binding (average dwell time τ_tb_) and diffusion in the nucleoplasm (with average duration τ_free_) as the molecule samples N_trials_ chromatin sites before reaching any one of the specific targets (Chen et al., 2014; Loffreda et al., 2017; Tatavosian et al., 2018). We found that τ_search_ of Mediator and the GTFs ranged between 8 s and 48 s (Figure 5B and see Methods), which is on the same order of magnitude as that of the yeast TF Ace1 (8 s) (Mehta et al., 2018) and generally ∼10-fold shorter than reported values for mammalian nuclear factors (Chen et al., 2014; Hansen et al., 2017; Tatavosian et al., 2018), likely reflecting the smaller yeast nucleus. Notably, target search predominantly took place in the nucleoplasm, with most GTFs undergoing 5-8 transient binding events (N_trials_) before stably engaging a target (Figure 5C). Intriguingly, Mediator and TFIID, with N_trials_>20, appeared to sample chromatin more frequently during PIC establishment. Of note, considering the vast number of potential sites represented by several thousand accessible NDRs genome-wide, the low numbers of N_trials_ sampled by PIC components suggest highly targeted recruitment with little non-specific interactions.

**Figure 5.**
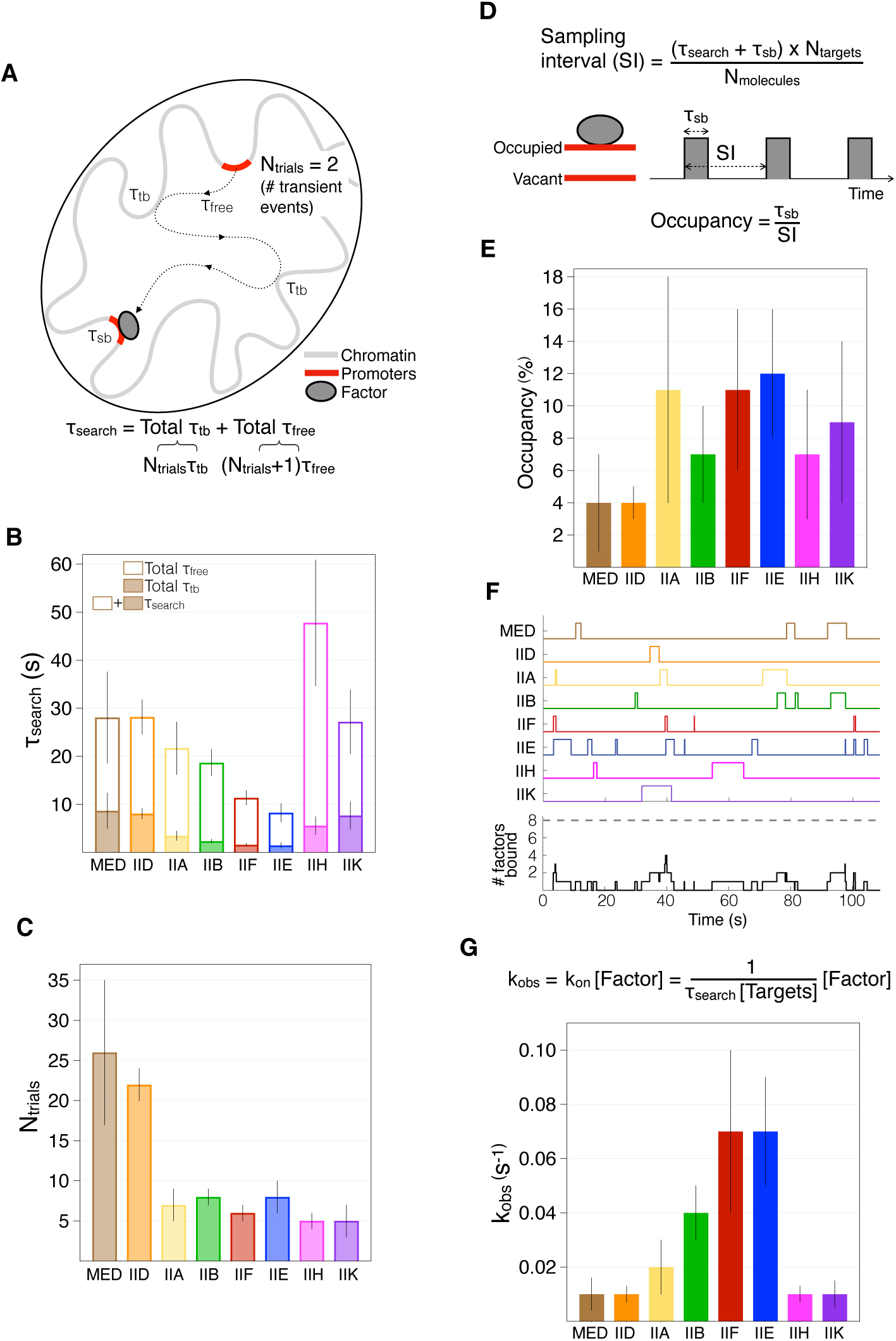
Target search kinetics and occupancy by PIC components. (A) Schematic of a single molecule’s trajectory between two promoter targets conceptualizing τ_sb_, τ_tb_, τ_free_ and τ_search_. The exemplary molecule diffuses in the nucleoplasm and samples N_trials_=2 nonspecific sites before encountering a target. The τ_search_ equation is indicated (see Methods). (B) Average total τ_free_ (white bars) and total τ_tb_ (colored bars) computed for each PIC component according to the equation in (g) (see Methods for τ_free_ and N_trials_ derivations). Errors were derived from three biological replicates in both fast and slow tracking. (C) Average N_trials_ of transient interactions between two specific binding events by each PIC component. Errors were derived from three biological replicates in both fast and slow tracking. (D) Schematic of unbound and bound states of an average promoter target conceptualizing the sampling interval (SI), τ_sb_ and temporal occupancy (O). The equations for calculating SI and O are indicated. The protein abundance of PIC components (N_molecules_ in molecules per cell) and number of PIC targets (N_targets_=6,000) were obtained from published results (see Methods). (E) Temporal occupancy (%) of each PIC component at an average promoter, calculated according to equation shown in (D). SI values are reported in Table S1. Errors were obtained from fast- and slow-tracking results (means ± SD) and documented # molecules/cell (median ± SD) (see Methods). (F) Association of PIC components at an average promoter assuming independent, uncoordinated binding, simulated over a 100 s time window based on SI and τ_sb_ values to represent occupancy levels close to those shown in (E). Bottom: cumulative plot showing the number of co-localized components, where ‘8’ is required for PIC establishment. Pol II and TBP occupancies are assumed based on dependencies established in Figures 2B and 2F. (G) Top: equation for calculating the observed association rate k_obs_ (s^-1^) of a pseudo-first order binding reaction between PIC components and chromatin targets, where the latter are considered mostly free at equilibrium due to low occupancy levels shown in (E). The concentrations of the PIC component ([Factor]) and chromatin targets ([Targets]) were derived from published data. Bottom: k_obs_ values obtained using the above equation. Errors were derived from τ_search_ results and SDs documented for cellular abundances of PIC components.

The % occupancy of the PIC machinery at a promoter target within a given time window is a critical metric of the gene’s potential for RNA synthesis. Target occupancy by a factor is a function of its target-search and dissociation kinetics as well as its abundance and the number of available targets (Figure 5D and see Methods) (Chen et al., 2014). Levels of Mediator and the GTFs range between 2,000 and 5,000 molecules per cell (Ho et al., 2018) and ∼6,000 PIC targets exist in budding yeast (Rhee and Pugh, 2012). These values indicate that PIC components each occupy an average promoter only 4-12% of the time (Figure 5E and Table 1). For example, the promoter target would host Mediator for 4 s (τ_sb_ from Figure 3C) once every 100 s, resulting in 4% occupancy (Figures 5D, equations). Based on these low individual occupancy levels, it would be highly improbable (p=10^-9^) for components to co-occupy a promoter to form the PIC without mechanisms to coordinate their recruitment (Figure 5F). Indeed, AA experiments demonstrated a strict recruitment hierarchy in live cells (Figure 2), indicating that occupancy by individual PIC components must temporally overlap for full PIC assembly and function. Consistent with such temporal convergence, we approximated similar observed association rates for Mediator, TFIID, IIH and IIK (k_obs_= 0.01 s^-1^, or one event every 100 s; Figure 5G). Intriguingly, TFIIB, IIE and IIF displayed ∼7-fold more frequent association, which may reflect a tendency to dynamically sample a target site, similar to the *in vitro* behavior of human TFIIB (Zhang et al., 2016), or perhaps re-association with the early elongation complex, as shown for TFIIF (Schweikhard et al., 2014; Zawel et al., 1995).

Together, these results reveal the dynamic nature of the target-search process by PIC components and provide kinetic parameters indicating that the PIC forms infrequently at an average yeast promoter. Moreover, the extrapolated association rates of individual components suggest that once nucleated, this process may occur with remarkable efficiency.

### Nucleosomes outcompete PIC at gene promoters

The essential remodeling enzyme RSC regulates the chromatin architecture of yeast NDRs. RSC inactivation leads to an upstream shift of the “+1” nucleosomes genome-wide and consequently occlusion of key DNA elements at a large subset of promoters (Ganguli et al., 2014; Hartley and Madhani, 2009; Kubik et al., 2018). To examine how nucleosome encroachment affects the recruitment and dynamics of PIC components, we inactivated RSC by AA depletion of the catalytic subunit Sth1 (Figures 6A and S6A). Fast-tracking experiments showed >50% reduction in chromatin binding by Pol II (Figures 6B, S6B and S6C), indicating a substantial transcriptional defect, in line with ChIP findings (Kubik et al., 2018). We also found ∼20-40% decreases in chromatin-bound fractions for Mediator, TFIIA, IIB, IIE, IIF, IIH and IIK (but not for TFIID), consistent with reduced TBP levels detected by ChIP at ∼30% of Pol II-transcribed genes, which were correlated with strongly shifted +1 nucleosomes following Sth1 AA (Kubik et al., 2018). (High levels of TBP binding to Pol I and Pol III-transcribed genes generally unaffected by RSC inactivation (S. Kubik and D. Shore, personal communication) may account for modestly decreased TBP binding detected in live cells.) These results suggest that the PIC is precluded from about a third of promoter targets after RSC inactivation, likely due to steric hindrance from the shifted +1 nucleosomes. The unexpected 50% increase in TFIID binding, shown for two different subunits: Taf1 (Figures 6B, S6B and S6C) and Taf2 (Figure S6D), suggests that RSC normally inhibits TFIID recruitment to chromatin, perhaps by directly competing for binding to the NDR and the +1 nucleosome (Brahma and Henikoff, 2018; Ramachandran et al., 2015).

**Figure 6.**
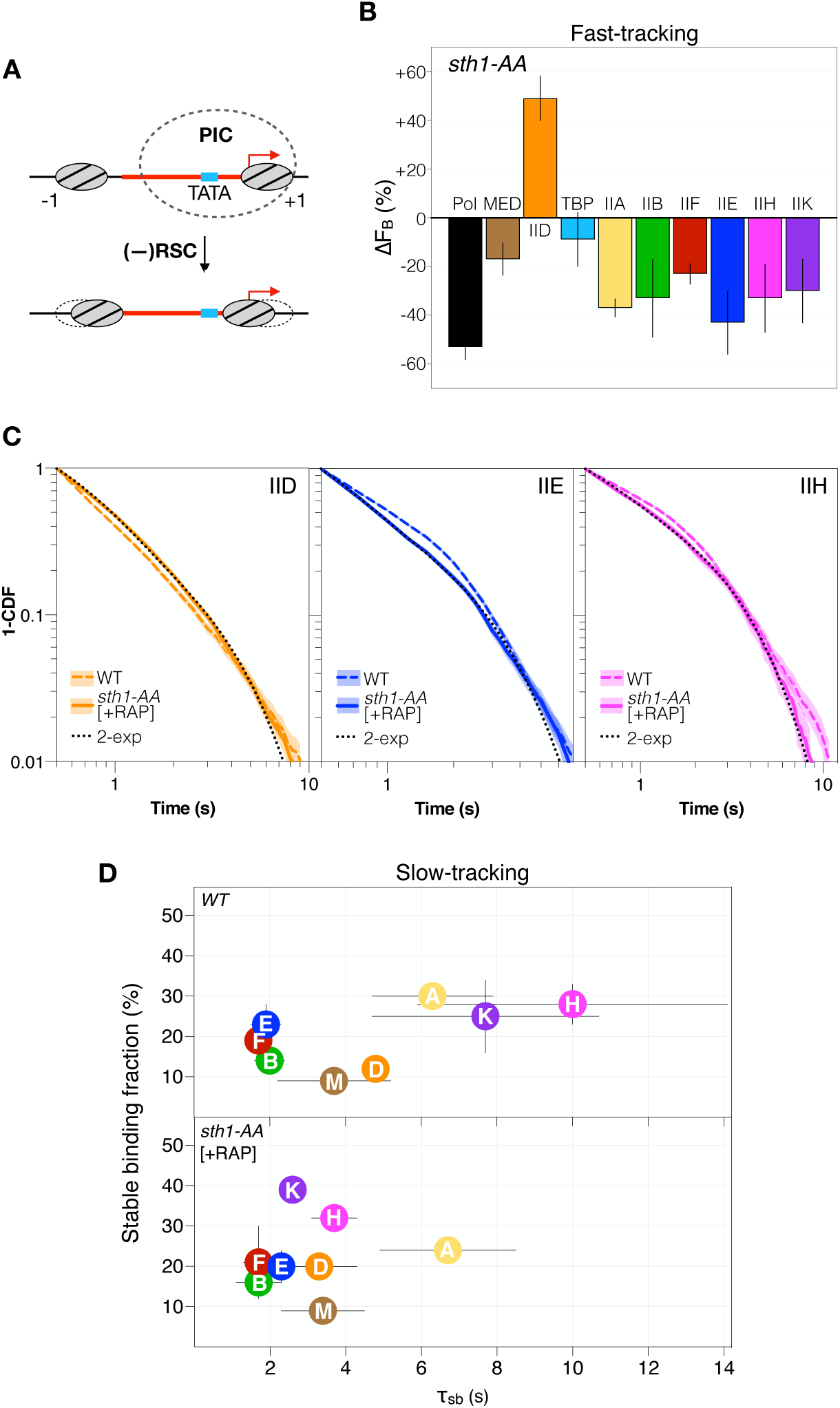
Nucleosome encroachment precludes PIC from many chromatin targets in live cells. (A) Top: Schematic of a yeast nucleosome-depleted region (NDR, red) featuring flanking nucleosomes (ovals) well-positioned relative to TBP-binding site (TATA) and the transcription start site (TSS, red arrow). Overlay of the PIC’s collective footprint (dashed oval) is informed by genome-wide mapping of components (Grünberg et al., 2016; Rhee and Pugh, 2012). Bottom: NDR “shrinkage” due to shifted flanking nucleosomes ensuing RSC inactivation. The upstream-shifted +1 nucleosome may prevent promoter access by the PIC. (B) ΔF_B_ after Sth1 depletion, assessed by fast tracking and shown as in Figure 2B. LogD histograms are reported in Figure S6B. Results were calculated using mean F_B_ ± SD, obtained by kinetic modeling (Figure S6C and Table S2), from three biological replicates of (-RAP) and (+RAP) conditions, and errors were propagated. (C) 1-CDF data of TFIID, IIE and IIH after Sth1 depletion, with corresponding wildtype data for comparison, shown as in Figure 4A (see Figures S7E and S7F for other components). [One *STH1 AA* sample without rapamycin was imaged to confirm wildtype-level dissociation kinetics (Figure S6E and Table S2).] (D) f_sb_ (%) and τ_tb_ (s) of PIC components in wildtype cells (top, *WT*) and after (bottom, +RAP) Sth1 depletion, shown as in Figure 4B. Double-exponential fit results are reported in Table S2.

To investigate whether RSC inactivation affects PIC dynamics at the large fraction of targets that remained accessible, we performed slow tracking and observed relatively unchanged dissociation kinetics (residence times τ_sb_) of Mediator, TFIID, IIA, IIB, IIE and IIF (Figures 6C, 6D, S6E and Table S1). Intriguingly, RSC inactivation resulted in shorter τ_sb_ observed for TFIIH and IIK, suggesting that RSC activity is also important for PIC functions post-assembly. Shortened temporal residence of TFIIH, which is tasked with promoter melting and TSS scanning (Murakami et al., 2015), is consistent with the correlation between TSS usage and changes in nucleosome positions upon RSC inactivation (Klein-Brill et al., 2019).

Taken together, our results indicate that RSC activity is important for PIC recruitment to a subset of promoter targets, and post-recruitment PIC functions at other targets. Furthermore, these findings demonstrate that the sensitivity of our SMT approach is amenable to dynamic changes following epigenetic perturbations in yeast.

## DISCUSSION

This study of PIC components in budding yeast reports multiple diffusive processes in the nucleoplasm and chromatin residence times under 5 seconds for most components, thus providing a limit for the physiological time window of PIC assembly and transcription initiation. Through conditional depletion experiments, we identify key factors coordinating nuclear exploration and chromatin recruitment, enabling individual components to converge, establish the PIC, initiate transcription and dissociate, all within a few seconds.

### Nuclear exploration by PIC components

All PIC components undergo subdiffusive processes in the yeast nucleus, with a tendency to explore 0.4 µm-0.5 µm radial regions. This behavior by the small GTFs TBP, TFIIA, IIB and IIE occurs in a fraction of the molecules, while other unbound molecules may search the entire nucleus (0.75 µm radius) (Figure 1F). Notably, the subdiffusive populations of TFIIB and IIE require the presence of Mediator (Figures 2C), and TBP-dependent subdiffusion by TFIIA (Figure 2A) likely occurs through interaction with TFIID (Kraemer et al., 2001). Furthermore, this exploration mode by TBP, TFIIA, IIB and IIE mirrors the diffusivity (Figure S2C), apparent confinement Rc (Figure 1F) and directional bias (Figure S2D) of Mediator and TFIID, suggesting that the large complexes constrain spatial exploration by the most diffusive GTFs. This process may collectively span the entire nucleus as individual molecules explore different subnuclear regions; alternately, it may be localized as molecules cluster in discrete locations. Intriguingly, ensemble fluorescence images indicate that the bulk of Mediator and TFIID co-occupy a subnuclear region consistent in size with the apparent confinement of single molecules (Rc∼0.4 µm) (Figure S7). Based on this information, we favor a scenario wherein Mediator and TFIID guide a fraction of TBP, TFIIA, IIB and IIE to explore a common subnuclear region while not excluding a more dispersive process contributing to PIC formation globally. Of further interest, the observed bulk distributions of Mediator and TFIID in yeast invite speculation of foci or condensates, which may be nucleated by sequence-specific TFs (Boija et al., 2018; Shrinivas et al., 2019), as shown for Mediator in mESCs (Cho et al., 2018), although we cannot discount volume exclusion, perhaps by the substantial nucleolus and other chromatin substructures, from giving rise to their compact organization (Woringer and Darzacq, 2018; Woringer et al., 2014). Such steric effects in the yeast nucleus would explain constrained diffusion of unbound large components—Mediator, TFIID, Pol II-TFIIF and TFIIH-IIK (>700 kDa)—compared to generally free diffusion of the small GTFs TBP, TFIIA, IIB and IIE (<150 kDa) (Figure 1F).

TFIIB and IIE subdiffusion and chromatin binding also requires the presence of Pol II in addition to Mediator (Figures 2E and 2F). We speculate that TFIIB and IIE could co-localize with a Pol II-Mediator complex, which has been biochemically purified from yeast nuclear extracts as a major form of Pol II (Rani et al., 2004). Furthermore, in the presence of Mediator and Pol II, interaction with nearby chromatin may constrain exploration by TFIIB and IIE to the local environment (Izeddin et al., 2014; McSwiggen et al., 2019). Such activity would be consistent with the relatively high chromatin association rates obtained for TFIIB and IIE, as well as IIF (Figure 5G). Importantly, in addition to TFIIB and IIE, Pol II is also required for recruitment of TFIIF, IIH and IIK. Thus, Mediator and Pol II constitute a critical bridge between chromatin targets and all enzymatic components of the PIC.

Taken together, we propose that Mediator and TFIID sequester enzymatic and promoter-recognizing components, respectively, to a shared subnuclear territory (Figure 7A), thereby establishing a dynamic network sustained by weak, rather than strong, interactions as components did not appreciably co-deplete in AA experiments (Figures S3A and S3G). This organization may facilitate PIC assembly by locally concentrating individual components, as Mediator and TFIID are required for basal transcription using physiological levels of factors but dispensable in a reaction with purified components (Baek et al., 2002). It may also enhance target-search efficiency by merging individual components in a reduced search space (Kent et al., 2020). Furthermore, the remarkably similar search and association kinetics of Mediator and TFIID in live cells (Figures 5B, 5C and 5G) complement biochemical, genetic and genomic evidence for their cooperative promoter engagement (Grünberg et al., 2016; Johnson and Carey, 2003; Johnson et al., 2002; Knoll et al., 2018). Therefore, Mediator and TFIID may serve to unite PIC components spatially in the nucleoplasm and temporally on chromatin targets. Our findings thus offer a dynamic, mechanistic perspective of their essential functions *in vivo* (Petrenko et al., 2017; Warfield et al., 2017). The model for spatial organization of PIC components also implicates a similar subnuclear distribution of highly transcribed genes in yeast, and future imaging studies may address colocalization of PIC components and active genes in more detail.

**Figure 7.**
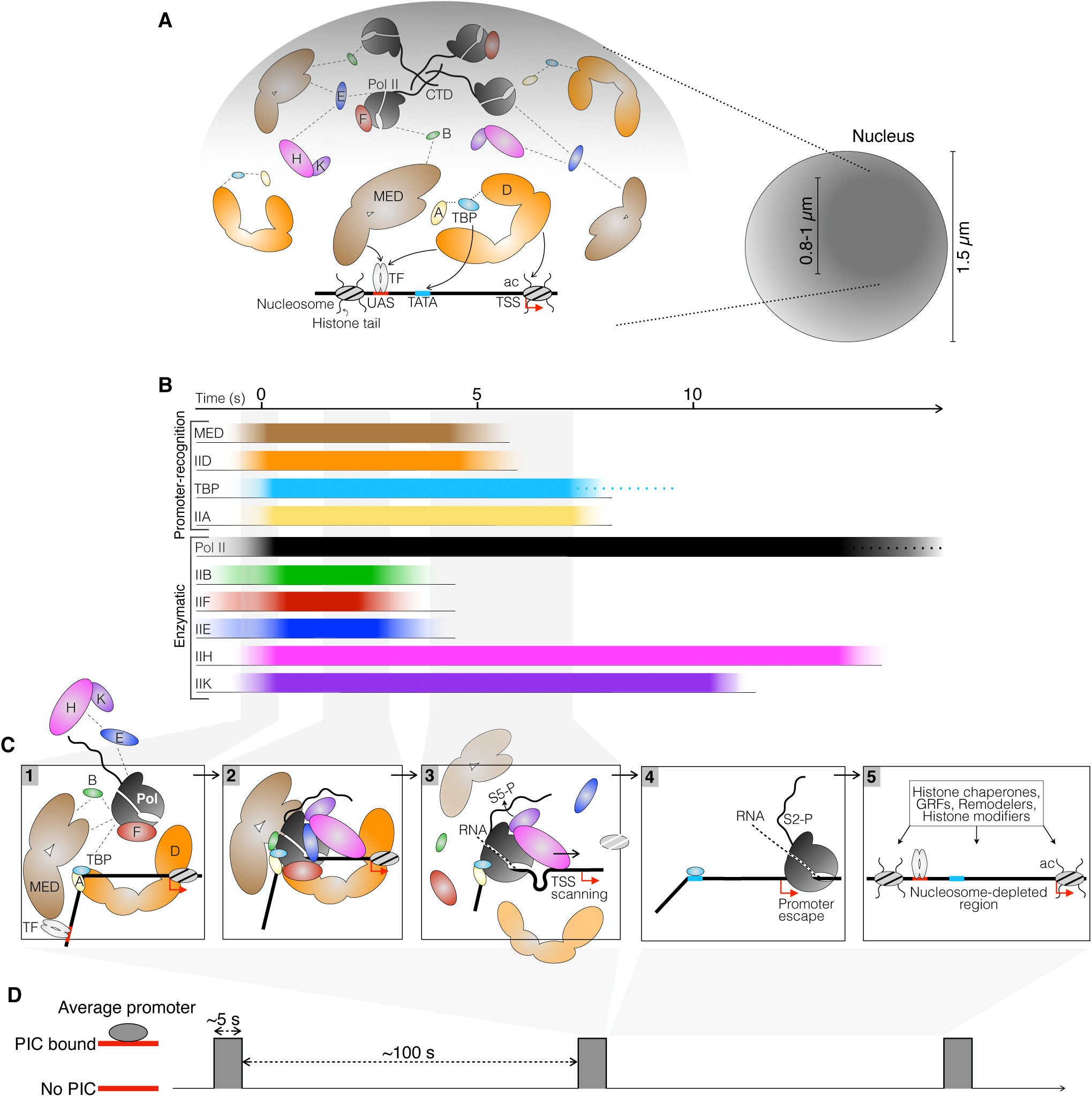
Spatial and temporal model of PIC establishment *in vivo*. (A) Left: A model for spatial clustering supported by multi-valent interactions (dashed lines) among promoter-recognizing PIC components TFIID, TBP and TFIIA as well as Mediator and enzymatic components. The presence of active genes in this region is implicated but only one stereotypical promoter is shown. Mediator and TFIID may be recruited by directly interacting with DNA elements, sequence-specific transcription factor(s) (TF), and the +1 nucleosome *via* acetylated histone tails (ac). Right: Schematic of a subnuclear environment explored by PIC components in a subdiffusive manner. The diameter of this region (dark shade) was estimated based on median Rc values (0.4-0.5 µm) (Figure 1F). (B) A temporal timescale of association and dissociation of PIC components at an average yeast promoter. Recruited components are shown to associate in a quasi-synchronous manner based on calculated k_obs_ (Figure 5G). Binding events are temporally scaled according to average τ_sb_ (Figure 3) and estimated for TBP and Pol II. (C) Sequential stages of PIC assembly and disassembly according to binding and dissociation of individual components. (D) A temporal schematic depicting sparse PIC establishment at an average promoter, based on the ∼5 s assembly window outlined in (D) and the association rate k_obs_ = 0.01 s^-1^ estimated for Mediator and TFIID (Figure 5G).

### A temporal model for PIC assembly and transcription initiation

We synthesize our SMT data with prior knowledge to propose a temporal sequence for *de-novo* PIC establishment at an average yeast promoter (Figures 7B). Mediator and TFIID survey the chromatin for NDRs, bound sequence-specific TFs, (Petrenko et al., 2016; Tuttle et al., 2018), GRFs (Layer and Weil, 2013; Papai et al., 2010) and local histone marks (Joo et al., 2017) (Figure 7A). Mediator and TFIID may engage the chromatin target cooperatively, perhaps through interactions with bound factors, such as Rap1 at ribosomal protein genes (Ansari et al., 2009; Layer and Weil, 2013), or through direct Mediator-TFIID interactions (Baek et al., 2006; Lim et al., 2007). TFIID and TFIIA orchestrate TBP engagement with TATA or TATA-like element (Patel et al., 2018; Zhang et al., 2016), effectively reconfiguring promoter architecture to distinguish it from bulk chromatin and other promoters (Figure 7C, Step 1). Mediator recruits Pol II, TFIIB, TFIIE and TFIIF to this reconfigured target, where TFIIE enables TFIIH/IIK recruitment to form a complete PIC (Compe et al., 2019; Maxon et al., 1994) (Figure 7C, Step 2). The coactivator SAGA, which is recruited by TFs to UASs (Baptista et al., 2017), might coordinate PIC assembly at a minor subset of promoters (Donczew et al., 2020), although its physical and functional overlap with Mediator is unclear.

TFIIK activity is associated with promoter disengagement by Pol II (i.e. promoter escape), Mediator (Wong et al., 2014) and TFIID (Knoll et al., 2020). In live cells, Kin28 depletion resulted in similar dissociation kinetics for Pol II, Mediator and TFIID (Figure 4B), suggesting that TFIIK-facilitated promoter clearance by these three components is kinetically coupled. Therefore, the normal ∼5 s average chromatin residence of Mediator and TFIID should encompass complete PIC assembly (i.e., after TFIIH/IIK recruitment) and key initiation events (including phosphorylation of Pol II CTD by TFIIK) (Figures 7B and 7C). Indeed, extrapolated association rates suggest that individual components may efficiently converge on a chromatin target, perhaps even in a synchronous manner (Figure 5C). Within the 5 s window, rapid dissociation of TFIIB and IIF may reflect TSS-scanning activity (Qiu et al., 2019) and synthesis of nascent RNA (Fujiwara et al., 2019) (Figure 7C, Step 3). Subsequent Pol II escape and entry into elongation may require mobilization or disassembly of the +1 nucleosome (Figure 7C, Step 4). Chromatin remodelers and regulators, including RSC and GRFs (Kubik et al., 2018), collaboratively reestablish the NDRs before the next initiation event (Figure 7C, Step 5).

We estimate that on average, this dynamic 5 s window for PIC activity occurs once every 100 seconds at a yeast promoter (Figure 7D), indicating that transcription initiation occupies a narrow temporal window at yeast NDRs. Dynamic traffic by RSC and other chromatin regulators, as well as the presence of fragile nucleosomes (Kubik et al., 2015), sub-nucleosome species (Brahma and Henikoff, 2018) or the Mot1-Ino80-NC2 complex (Xue et al., 2017) may collectively finetune NDR access by the PIC. Importantly, considering the median nascent transcription rate of 7 mRNAs/hour among genes expressed during exponential growth in glucose (Pelechano et al., 2010), our global estimate of PIC association kinetics—up to 36 PICs/hour—suggests that most PICs either fail to initiate, or initiate without completing the transcription cycle. Such spurious initiation from promoters has been implicated as a major source of pervasive cryptic transcripts (Neil et al., 2009; Xu et al., 2009). On the other hand, inducible genes in yeast exhibit transcriptional bursts ranging between once every ∼8 s (Mehta et al., 2018) and 4 minutes (Donovan et al., 2019), implicating additional mechanisms regulating PIC kinetics that may be revealed in future by SMT in the local neighborhood of specific genes.

In summary, our study deconstructs global yeast transcription initiation in physiological space and time, revealing specific coordination on and off chromatin that enables efficient complex assembly. Our findings provide an average temporal guide to future investigations of gene-specific transcription kinetics and offer insights relevant to other processes involving establishment of multi-component machineries at sparse chromatin targets.

## ACKNOWLEDGMENTS

We dedicate this work to the memory of Maxime Dahan, former leader of the Transcription Imaging Consortium (TIC) at the HHMI-Janelia Research Campus. We thank Anders Hansen, Maxime Woringer and Xavier Darzacq for assistance with Spot-On and angle distribution analyses; Slawomir Kubik and David Shore for sharing unpublished results; Diana Stavreva, David Garcia, Arpita Upadhyaya and Gordon Hager for input on statistical assessment of residence time data; Lu Bai for technical advice on FRAP in yeast; Xiaojun Ren for SMT discussions; Erin Pryce and the Integrated Imaging Center for training on the LSM 800 for FRAP experiments; Matt Hurlock and Yumi Kim for sharing resources and assistance with deconvolution microscopy, and Gisela Storz, Jessica Liu, Rogelio Hernandez-Lopez and Wu Lab members for comments. This study was supported by HHMI funding to the TIC (C.W., T.L., Q.Z., and L.L.), the Damon Runyon Cancer Research Foundation (V.Q.N.), a Johns Hopkins Bloomberg Distinguished Professorship (C.W.) and the National Institute of Health grant GM132290-01 (C.W.).

## AUTHOR CONTRIBUTIONS

V.Q.N. performed all genetic and imaging experiments with support from A.R., J.W., G.M., K.Y.L. and V.J, and all data analysis using R functions created by S.L., X.T. and Y.H.L.

L.D.L. and Q.Z. synthesized JF^552/646^. T.L. provided overall guidance on SMT data analysis, derived mathematical calculations of search kinetics, and generated the MATLAB simulations. V.Q.N. and C.W. designed the study and wrote the paper with input from all authors.

## METHODS

### Yeast strain construction

Proteins were chromosomally tagged in haploid W303 *Saccharomyces cerevisiae* by standard high-efficiency transformation and homologous recombination of PCR-amplified DNA (Gietz and Schiestl, 2007). All yeast strains harbored *pdr5*Δ to enhance retention of fluorescent ligands (Ball et al., 2016). To tag the C terminus, the *HALOTAG* sequence (Promega) was cloned into the pBluescript SK (-) vector upstream of a NatMX (nourseothricin) cassette, generating plasmid pBS-SK-Halo-NatMX. PCR-amplified DNAs for yeast transformation contained *HALO-NATMX* with a 5’ sequence coding for a SG_4_ amino-acid linker. To tag the N termini of TBP and Toa1, the *HALOTAG* sequence was cloned between the PacI and AscI restriction sites on pFA6a-TRP1-pGAL1-HBH (Booher and Kaiser, 2008), generating plasmid pFA6a-TRP1-pGAL1-HALO. PCR-amplified DNAs for yeast transformation contained sequences coding for a GSG_4_ (TBP) and (G_4_S)_2_ (Toa1) linker. Tagging was carried out according to the strategy outlined by Booher & Kaiser for essential proteins (Booher and Kaiser, 2008).

For Anchor-Away (AA) experiments, *pdr5*Δ was implemented in strain HHY221 (Haruki et al., 2008). Then, the *FRB-GFP-KANMX* sequence was synthesized from plasmid pFA6a-FRB-GFP-kanMX to target *RPB1, KIN28,* and *STH1* at the 3’ ends. Strains expressing Rpb1-FRB-GFP, Kin28-FRB-GFP, and Sth1-FRB-GFP were confirmed for rapamycin-dependent nuclear depletion my imaging GFP fluorescence. To create a strain expressing FRB-GFP-TBP for TBP AA, N-terminal tagging was carried out as described with PCR-amplified DNA containing *FRB-GFP* and a sequence coding for G_4_SG_3_SG_4_ linker (from plasmid pFA6a-TRP1-pGAL1-FRB-GFP). Subsequently, PIC components were HaloTagged in AA strains as described.

To generate yeast strain expressing Halo-H2B, we replaced the *SNAP* sequence in *HTA1-SNAP-HTB1* on pEL458 (gift from Ed Luk) with *HALOTAG* sequence encoding a GAAA linker to the N-terminus of H2B. The resulting plasmid (*HTA1-HALO-HTB1*) was shuffled (Hirschhorn et al., 1995) into the FY406 strain (gift from Fred Winston). Sole-source Halo-H2B was expressed under control of its endogenous promoter.

To generate yeast strain expressing nuclear HaloTag, a sequence coding for a bipartite SV40 nuclear localization signal (NLS) (N-KRTADGSEFESPKKKRKV-C) was fused to the 5’ end of *HALOTAG* and inserted into the pRS416 vector to generate pRS416-NLS-HALO. The NLS sequence was acquired from plasmid pAC1056 (gift from Anita Corbett).

### Yeast cell growth and spot test

Cells were grown at 30°C in YAPD. Serial 6-fold dilutions in YAPD were prepared from OD_600_ = 1.0 cultures. Dilution series were spotted on YAPD plates and incubated in the dark at room temperature, 30°C, and 38°C for two days to assess the heat-shock response. We also spotted W303 strain harboring no Halo fusions and *htz1*Δ strain as negative and positive controls for growth defect, respectively.

### Live-cell fluorescence imaging

#### Sample preparation

Cells were grown at 30°C in CSM in the presence of 3-30 nM (for fast tracking) or 1-10 nM (for slow tracking) JF^552^ to label Halo fusions. Mid-log cultures (OD_600_∼0.6-0.8) were harvested and washed with ∼1 mL CSM five times. For AA experiments, rapamycin (LC Laboratories), dissolved in DMSO, was added to the growing culture (OD=0.6) to a final 1 µg/mL concentration 1 hour before harvesting. All washes were carried out with CSM + rapamycin (1 µg/mL) medium. For control AA experiments, DMSO was added to the culture and wash medium.

A coverslip (#1.5, ø 25 mm, Electron Microscopy Services) was heat-treated, coated with Concanvalin A (0.5 mg/mL), and assembled in a metal Attofluor chamber (ø 35 mm, Invitrogen). Washed cells (1 mL) were added and allowed to attach to the coverslip for 2 minutes, followed by gentle rinses with fresh wash medium to achieve a monolayer of cells. Final sample contains 1 mL of medium in the chamber.

#### Single-molecule imaging

Imaging was carried out at room temperature on an Axio Observer Z1 microscope (Zeiss, Germany) equipped with an α-Plan-Apochromat 150x/1.35 glycerin-immersion objective (ZEISS, Germany). To excite JF^552^, we used CL555-100 555 nm laser (CrystaLaser, Reno, NV) and a filter cube containing a 561 nm BrightLine single-edge beamsplitter and a 612/69 nm BrightLine single-band bandpass emission filter (Semrock, Rochester, NY). Images were acquired by a C9100-13 EM-CCD camera (Hamamatsu Photonics, Japan) featuring 512x512 pixels with 16 µm pixel size, operating at ∼-80°C (forced-air cooling) and EM gain 1200x. The pixel size of recorded images is 107 nm. A 750 nm blocking edge BrightLine multiphoton short-pass emission filter and a 405/488/561/635 nm StopLine quad-notch filter (Semrock, Rochester, NY) were placed in front of the camera. We used the ZEN imaging software (ZEISS, Germany) and HCImage (Hamamatsu Photonics, Japan) to operate the microscope and camera, respectively. Excitation laser was triggered by TTL exposure output signal from the camera.

For fast tracking, we excited the sample with ∼1 kW/cm^2^ continuous laser and imaged a 128x128 pixel field of view (containing ∼5 single cells) for ∼1.5 minutes at 10 millisecond camera integration time. For slow tracking, we used lower excitation power (∼50 W/ cm^2^) and imaged a 256x256 pixel field for ∼3 minutes at 250 ms/frame. For time-lapse imaging (Gebhardt et al., 2013) of H2B, Rpb1 and TBP, we modified the slow-tracking regime to alternate 250 ms excitation and 250 ms or 500 ms dark time and imaged each field of view for ∼3 minutes at 500 ms or 750 ms/frame, respectively.

## QUANTIFICATION AND STATISTICAL ANALYSIS

### Localizing and tracking single molecules

Raw movies were pre-processed in Fiji (Schindelin et al., 2012) to bypass bleaching of initial nuclear fluorescence (first ∼1,000 and 150 frames for fast and slow tracking, respectively) and generate a substack (5,000-6,000 and 600-1,000 frames for fast and slow-tracking files, respectively) in which each image contained sparse single molecules (<2 molecules per nucleus). We used a maximum-intensity Z projection of each file to locate and manually window nuclei to create a binary mask. Substacks were saved as 16-bit TIFFs for subsequent analysis.

To localize single molecules, we used DiaTrack v3.05 (Vallotton and Olivier, 2013) at the following settings: high precision (HWHM=1 pixel), remove blur 0.1, remove dim 50-100. Tracking was performed with 6 pixel (∼0.65 µm) maximum allowance between consecutive localizations. This cut-off was informed by the smooth tail of the frequency histogram of displacements (Figure S1E). In some cases, such as free HaloTag, 8 pixels were allowed to achieve this feature. Localization and tracking results were saved as MATLAB files for subsequent analyses.

Slow-tracking data were similarly processed, with 2-pixel maximum allowance between consecutive localizations based on H2B displacement observed under the same imaging regime (Figures S4A and S4B).

### Analyzing fast-tracking data

We used the Sojourner package (https://github.com/sheng-liu/sojourner) to analyze tracking results. Trajectories containing < 3 localizations (2 displacements) were discarded to reduce false detections. Binary masks generated during pre-processing were applied to select nuclear trajectories. Average length of selected data was 10-12 displacements (median = 5-7). To determine the diffusion coefficient (D), we performed linear fitting between Δt 20 – 50 ms of the MSD computed for trajectories containing ≥ 5 displacements. The slopes of the lines, subject to R^2^≥ 0.8 criterion, were divided by 4 to obtain D (Mazza et al., 2012). We then imported log_10_D values to Prism (version 8.00 for Mac, GraphPad Software, La Jolla, California, USA, www.graphpad.com) and plotted the frequency histogram with 0.2 µm^2^/s binning of logD values. We used the default “Sum of two Gaussians” and a custom “Sum of three Gaussians” function to fit the logD distributions without hard constraints. We did not quantify the resolved subpopulations due to limits of the MSD-based analysis, including the trajectory-length selection and unreliable linear fits of MSDs from very stably-bound molecules (data not shown), which omitted up to 50% of the data. We used this approach to inform the number of resolvable dynamic states for each factor. To perform quantitation of these states, we carried out two- or three-state kinetic modeling of displacements obtained from all trajectories (Hansen et al., 2018). We used the Spot-On web interface (https://spoton.berkeley.edu/) with the following settings: bin width = 0.01 µm, number of timepoints = 5, jumps to consider = 6, max jump = 1-1.2 µm and Z correction with dZ = 0.6 µm. Model fit was performed on the CDF of displacements for 3 iterations. The localization error, obtained from fitting the data, was ∼25-40 nm. Data for HaloTag and chromatin-binding mutants where the average trajectories were shorter (4-6 displacements) were fitted to 3-4 timepoints and with 0.35 nm localization error constraint.

### Subclassifying trajectories and MSD analysis

We separated trajectories from distinct subpopulations according to logD values within one standard deviation of the mean. After this process, two or three subsets were generated for each PIC component and H2B, without significant overlap (Figure S2A). We then computed the average MSD for each subset and performed power-law fit MSD=B(Δt)^α^ between Δt = 10 ms and 100 ms (Figure S2B). A power-law behavior where α<1 indicates sub-diffusion (higher confinement ∼ smaller α) and B is directly related to diffusion coefficient (Izeddin et al., 2014). The average MSD profiles of chromatin-free subpopulations were consistent with free diffusion in a confined space, where the MSD plateau is related to the diameter of the confinement (Dion and Gasser, 2013). The theoretical MSD plateau for free diffusion in the haploid yeast nucleus (diameter∼1.5 µm) is 0.45 µm (Dion and Gasser, 2013) (Figure S2B, dotted line).

To calculate the apparent radius of confinement (Rc) for individual trajectories, we fit each MSD curve with the power-law function as an initial selection for subdiffusive molecules (α <1). MSD curves from these molecules were then fit to the following confined diffusion model (Lerner et al., 2020; Wieser and Schütz, 2008),

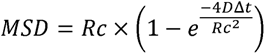

### Displacement angle analysis

We calculated the angles formed to consecutive displacements of single molecules according to Hansen *et al*. (Hansen et al., 2020). The analysis was performed on the subclassified intermediate and unbound populations of Mediator, TFIID, TBP, TFIIA, IIB and IIE. Backward bias of molecule movement, i.e. 180°±30° / 0°±30° > 1, indicates potential confinement.

### Calculating change in chromatin-bound fraction after AA

For AA experiments, we calculated the change in chromatin-bound fraction F_B_ (%) as follows,

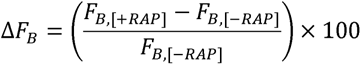

F_B,[+RAP]_ was determined for three biological replicates. For *rpb1-AA*, *tbp-AA*, and *kin28-AA*, we obtained F_B,[-RAP]_ for one biological replicate to confirm wildtype-level association frequencies (Table S2). Then, we used F_B,WT_ to calculate ΔF_B_. For *sth1-AA*, F_B,[-RAP]_ was determined for three biological replicates. All variables applied were means ± s.d. from biological replicates and the error for ΔF_B_ was obtained through propagation.

### Analyzing slow-tracking data

Tracking results from DiaTrack were processed in Sojourner to determine the apparent dwell times (temporal lengths of trajectories). The cumulative frequency distribution (1-CDF) of dwell times was fitted with the double-exponential decay function, where f_sb_ and k_sb_ represent the frequency and apparent dissociation rate constant of stable binding, respectively, and k_tb_ represents the apparent dissociation rate constant of transient binding (Mazza et al., 2012),

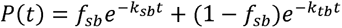

We used the bootstrap method (Efron and Tibshirani, 1986) to generate 100 resampled datasets and calculate 95% confidence interval (CI). We then computed 1-CDF and performed fitting for each dataset. The mean values for f_sb_, k_sb_, and k_tb_ were taken, with their SDs providing an assessment of fit quality.

We adapted the approach described by Hansen *et. al.* (Hansen et al., 2017) to correct the apparent dissociation rates. First, we imaged chromosomal H2B under the same regime, with the assumption that its decay kinetics measures limits imposed by photobleaching and chromatin motions. The 1-CDF for H2B was fitted as described (Figure S4C), and its k_sb_ was used for correction as follows,

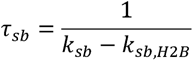

### Calculating search kinetics

Our derivations represent a hybrid of approaches described by Tatavosian *et. al.* (Tatavosian et al., 2018) and Loffreda *et. al.* (Loffreda et al., 2017). There are N_ns_ nonspecific and N_s_ specific binding sites in the genome, associated with transient (average lifetime τ_tb_) and stable (average lifetime τ_sb_) binding events, respectively. A molecule’s typical trajectory cycle is composed of N_trials_ nonspecific interactions interspersed with free diffusion (lasting on average τ_sb_), followed by a specific binding event. Assuming equal on-rates at all genomic sites, the molecule samples all sites at random and the average search time between two consecutive specific binding events is,

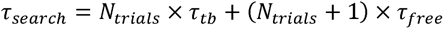

Assuming equal accessibility and introducing the nonspecific-to-specific site ratio as r_s_ = N_ns_/N_s_,

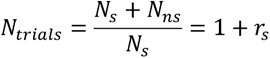

Thus,

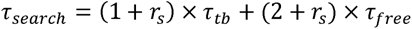

To obtain r_s_, we considered two scenarios underlying detection of binding events during slow tracking.

(a) *Blinking-limited*: molecules fluoresce in a bound state with probability P_s_ of occurring at a specific site, which is equal to the fraction of stable binding events obtained from slow tracking. Ps is proportional to the fraction of time the molecule spends at specific sites relative to the overall time it spends bound to chromatin.

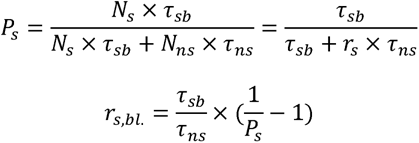

(b) *Diffusion-limited*: molecules fluoresce in a diffusive state and are motion-blurred until they engage a binding site in focus. Here, P_s_ is proportional to the number of specific sites relative to all sites,

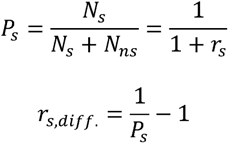

These physical processes can happen in the cell coincidentally and likely represent two extremes of a spectrum of behaviors single molecules can exhibit. We reasoned that the relative likelihood of detecting a blinking-limited binding event by slow tracking is proportional to the global fraction of bound molecules (F_B_). Therefore, we computed a weighted average value for r_s_ as follows,

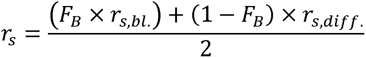

To obtain τ_free_, we considered the chromatin-bound frequency F_B_, obtained by fast tracking, as proportional to the fraction of time the molecule is bound at specific and non-specific chromatin sites,

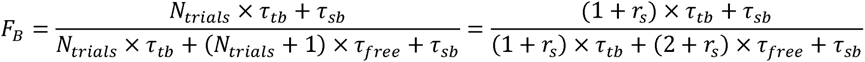

Therefore,

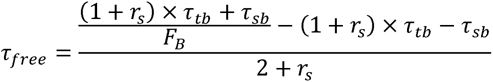

### Calculating target occupancy

We used the approach described by Chen *et. al.* (Chen et al., 2014) to calculate the average temporal occupancy of each PIC component at an average promoter target. First, we estimated the sampling interval SI, which represents the average time between two consecutive binding events at a specific site,

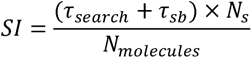

We considered N_s_=6,000 PIC targets based on published ChIP-exo results (Rhee and Pugh, 2012). Values (median±s.d) for cellular abundance (N_s_, molecules/cell) were obtained from the *Saccharomyces* Genome Database (SGD) (Cherry et al., 1998)—Med14: 1,989±849; Taf1: 1,633±434; Toa1: 2,599±1,754; Sua7: 4,262±1,425; Tfg1: 4,780±2,343; Tfa1: 3,514±437; Ssl2: 2,137±600; Kin28: 2,151±704.

The average target occupancy (O) is,

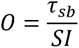

### Simulating target occupancy

We used experimentally determined τ_sb_ and calculated SI to simulate occupancy of each PIC component at an average promoter according to the null model of all PIC factors binding independently. For each factor, we simulated a sequence of on/off intervals during which the promoter was respectively occupied and unoccupied. During the simulation, the duration of individual on/off intervals were drawn at random from exponential distributions with respective average *τ_sb_* and SI - *τ_sb_*. We chose the exponential distribution form because it corresponds to the simplest possible biochemical scenario of a single rate-limiting step driving the lifetime of each state. Once we independently simulated on and off states for each of the PIC components at the promoter, we scanned through the time traces to calculate the number of factors bound at each given time. Simulations were performed in MATLAB.

### Fluorescence Recovery after Photobleaching (FRAP)

Haploid strains expressing Halo-H2B, Rpb1-Halo, and Halo-TBP were grown and imaging samples prepared as described. We labeled Halo fusions with 20 nM JF^646^ (Grimm et al., 2015). FRAP experiments were performed at room temperature on an LSM 800 confocal microscope with Airyscan and a Plan-Apochromat 63x/1.40 oil-immersion objective (ZEISS, Germany). Microscope control and data acquisition were performed in Zen (ZEISS, Germany). Two frames of each cell were acquired before photobleaching. Then, a 640 nm laser was used at 100% power for ∼10-30 to bleach ∼50% of the nuclear region, and the cell was excited with 0.2% laser power and imaged at 1 s interval for about 2 minutes. For each sample, FRAP was performed on 20-25 cells.

Data were analyzed in Fiji. We used the StackReg plugin (Thevenaz et al., 1998) to correct for drift when applicable. For each frame, mean intensities were determined and background-subtracted for a reference area of unbleached fluorescence (REF) and the bleached area (BL). BL intensity relative to the REF (BL/REF) was normalized against pre-bleach BL/REF to account for the maximum intensity achievable by fluorescence recovery. Data from 15-20 cells were averaged to generate a single recovery curve for “Exponential Recovery” fit up to 30 s due to significant movement of cells past this time point. The fit function was *y* = *a* x (*a* - *e*^-*bx*^)+*c*, where a represents the fraction of slow recovery and b the corresponding rate, and 1/b the estimated τ.

### Deconvolution microscopy

Yeast cells were grown to mid-log (JF^646^ was added to label Taf1-Halo), harvested and fixed using 4% formaldehyde (EMS) in PBS for 15 minutes at room temperature. Fixed cells were washed with PBS and spread between a coverslip and a standard glass slide. The edges of the imaging sample were sealed with clear nail polish.

Imaging was performed at room temperature on a DeltaVision Elite system (GE Healthcare) equipped with an Olympus 100 × 1.4 NA oil-immersion objective and a scientific complementary metal-oxide semiconductor camera (PCO). For each field of view, GFP and JF^646^ fluorescence image stacks were acquired sequentially at 0.2-µm intervals. Individual stacks were subjected to signal ratio enhancement and 20 cycles of iterative deconvolution.

Projection images were generated by the Volume Viewer tool using SoftWoRx suite (GE Healthcare). The resulting images were colored and merged in ImageJ.

## SUPPLEMENTAL INFORMATION

**Figure S1.**
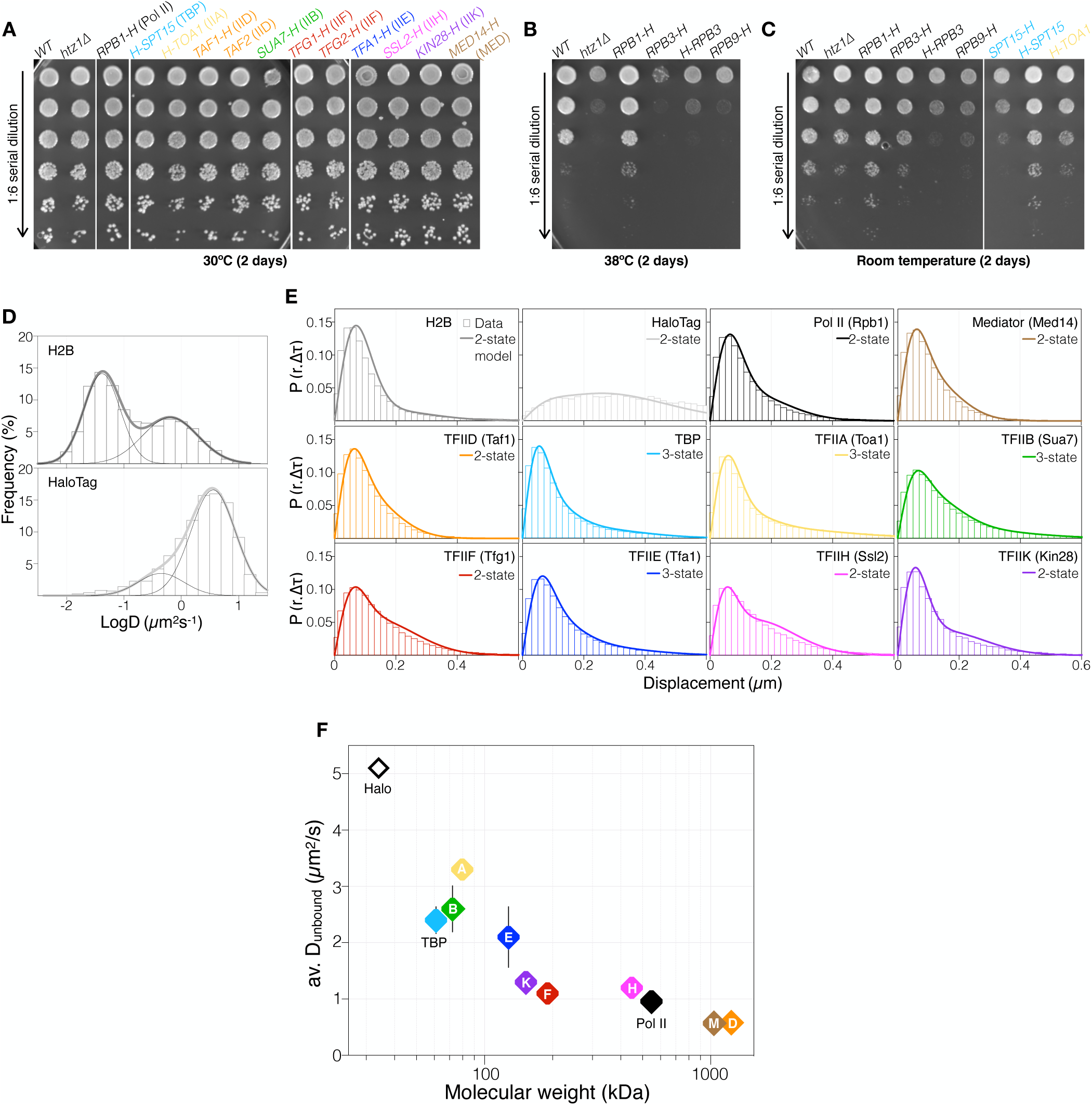
Global diffusivities of individual PIC components, Related to Figure 1. (A) Growth phenotype assay at 30°C confirming functionality of HaloTag (H) fusions in individual yeast strains. *htz1*Δ strain serves as a control for growth defects under certain conditions. (B) and (C) C- and N-terminal H fusions of Rpb1 and TBP, respectively, exhibiting wildtype-level growth at 38° (B) or room temperature (C) compared to other genetic permutations. (D) LogD histograms (bars), two-Gaussian fits (thick curves) and subpopulations (thin black curves) of H2B (top) and free nuclear Halotag (bottom). Histograms contain data from three (H2B) or two (HaloTag) biological replicates. (E) Displacement (Δt=10 ms) distributions (bars) and corresponding two- or three-state kinetic models (lines). Modeling results are found in Table S1. Histograms contain data from two (HaloTag) or three (others) biological replicates. (F) Correlation between D_free_ (from kinetic modeling) and theoretical molecular weight of each PIC component (log_10_). D_free_ values, reported in Table S1, are means ± SD from two (HaloTag) or three (others) biological replicates.

**Figure S2.**
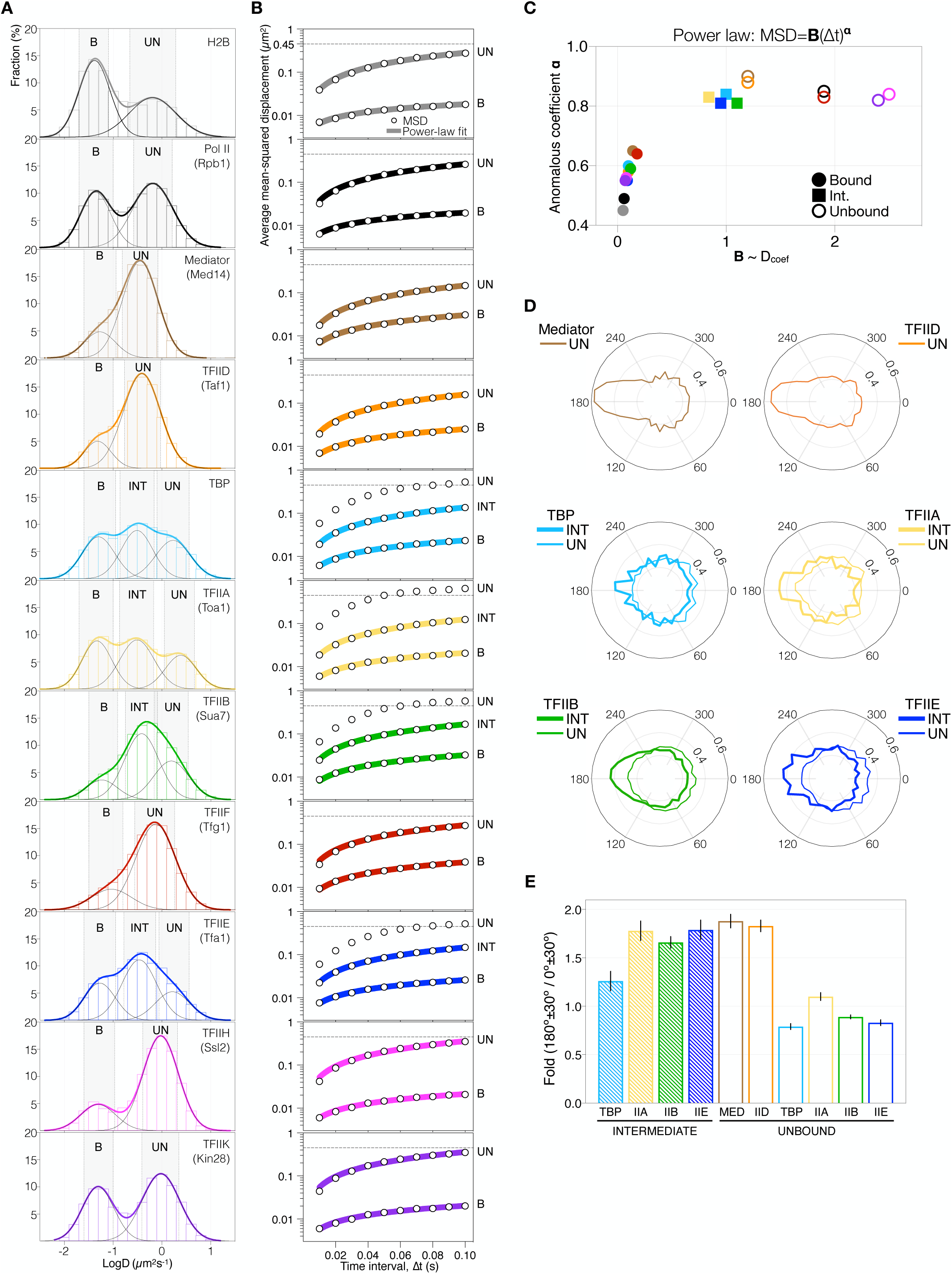
Sub-classification of trajectories and mean-squared displacement (MSD) analysis of distinct dynamic behaviors, Related to Figure 1 (A) Fitted logD histograms as in Figure 1C. Trajectories exhibiting D values within mean±1 SD of respective subpopulation (shaded bars) were selected as a subclass for MSD analysis. B, chromatin-bound. UN, unbound. INT, intermediate. (B) Average MSD (log_10_) computed for each subpopulation and corresponding power-law fit (solid line) of CB and INT. Average MSD plots of free diffusion plateau near the expected MSD ∼ 0.45 µm^2^ (dashed line, see Methods) for such behavior in the haploid yeast nucleus (0.75 µm radius). (C) Power-law fit (equation indicated on top) results showing the anomalous coefficient α, where α<1 indicates a subdiffusive behavior, and the constant B, which is proportional to the average D_coef_. (D) Angle analysis for unbound (UN) and intermediate (INT) populations of Mediator, TFIID, TBP, TFIIA, IIB and IIE. Normalized angle distributions were obtained using the vbSPT package (Hansen et al., 2020) (see Methods). (E) The probabilities of backward (large 180°±30° angles) relative to forward (small 0°±30° angles) movement, computed based on the angle distributions in (D) (see Methods).

**Figure S3.**
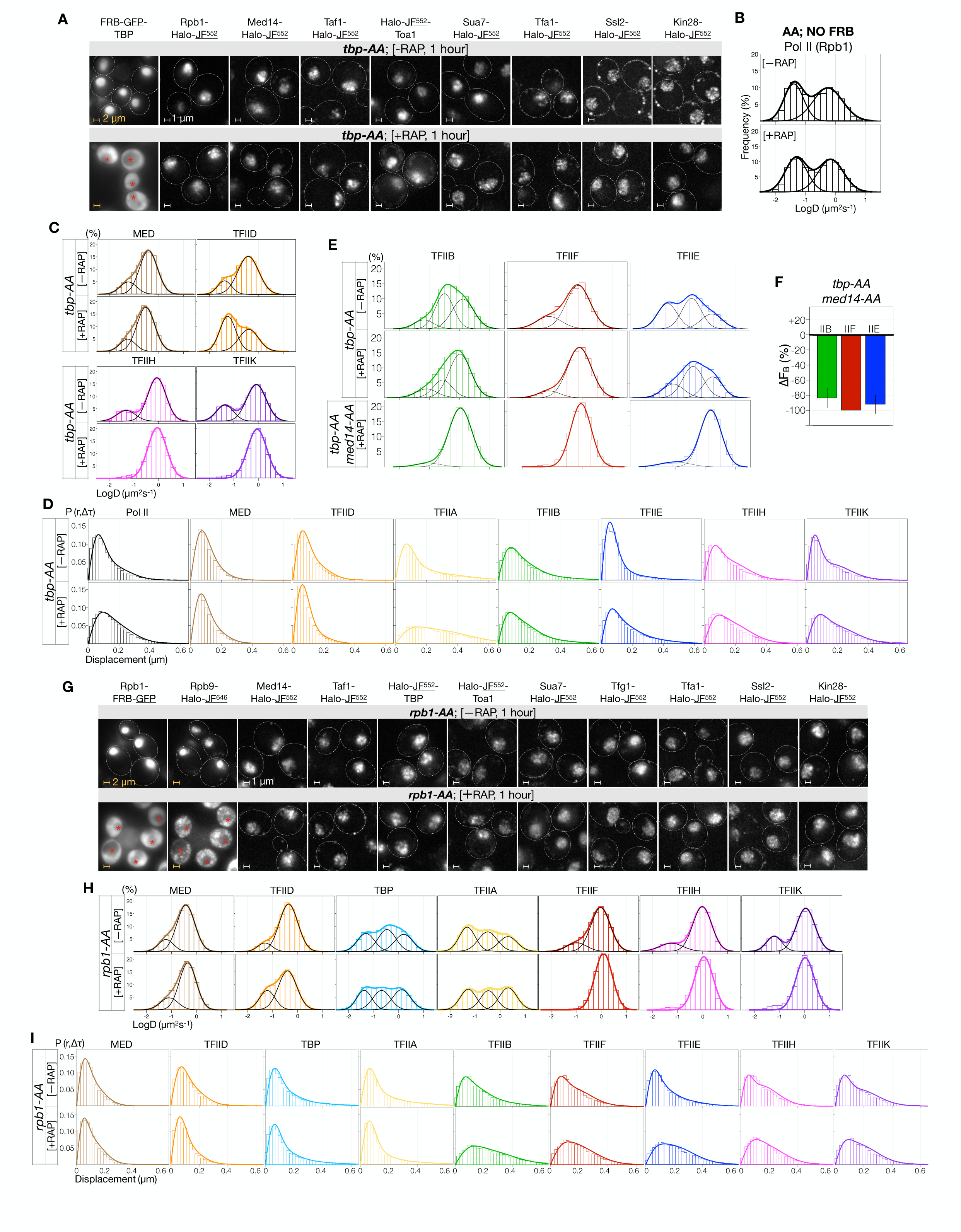
Effect of TBP and Pol II depletion on dynamics of PIC components, Related to Figure 2. (A) Z-projection images showing GFP (TBP) and Halo-JF (others) fluorescence in *tbp-AA* cells before (-RAP, top) and after rapamycin addition (+RAP, bottom). Cell borders are indicated as dashed ovals. Growing cells were treated with DMSO (-RAP) or 1 µg/mL rapamycin (+RAP) for 1 hour before imaging. Imaged fluorescence is underlined. *, vacuole. Scale bar: 2 µm (yellow) and 1 µm (white). (B) LogD histograms of Rpb1-Halo in a control Anchor-Away (*AA*) strain harboring no FRB-tagged protein. Rapamycin treatment as described in (A) had little effect on global Pol II dynamics. (C) LogD histograms of PIC components in *tbp-AA* cells before (-RAP, top) and after (+RAP, bottom) Pol II depletion, shown as in Figure 2A. Cells were treated as described in (A) before fast tracking for ∼2 hours. Histograms contain data from one (-RAP) or three (+RAP) biological replicates. (D) Kinetic modeling for data in (C) and Figures 2A and 2C, shown as in Figure S1E. Results are reported in Table S2. (E) LogD histograms of TFIIB, IIF and IIE before (-RAP) and after (+RAP) TBP depletion (*tbp-AA*), as well as those obtained after simultaneous TBP/Med14 depletion (*tbp-AA*; *med14-AA*). Data are shown as in Figure 2C. (F) F_B_ changes relative to wildtype after double TBP/Med14 depletion, computed and shown as in Figure 2B. (G) Z-projection images of PIC components in *rpb1-AA* cells, shown as in (A). (H) LogD histograms of PIC components in *rpb1-AA* cells, shown as in (C). (I) Kinetic modeling of data in (B) and Figure 2E. Results are reported in Table S2.

**Figure S4.**
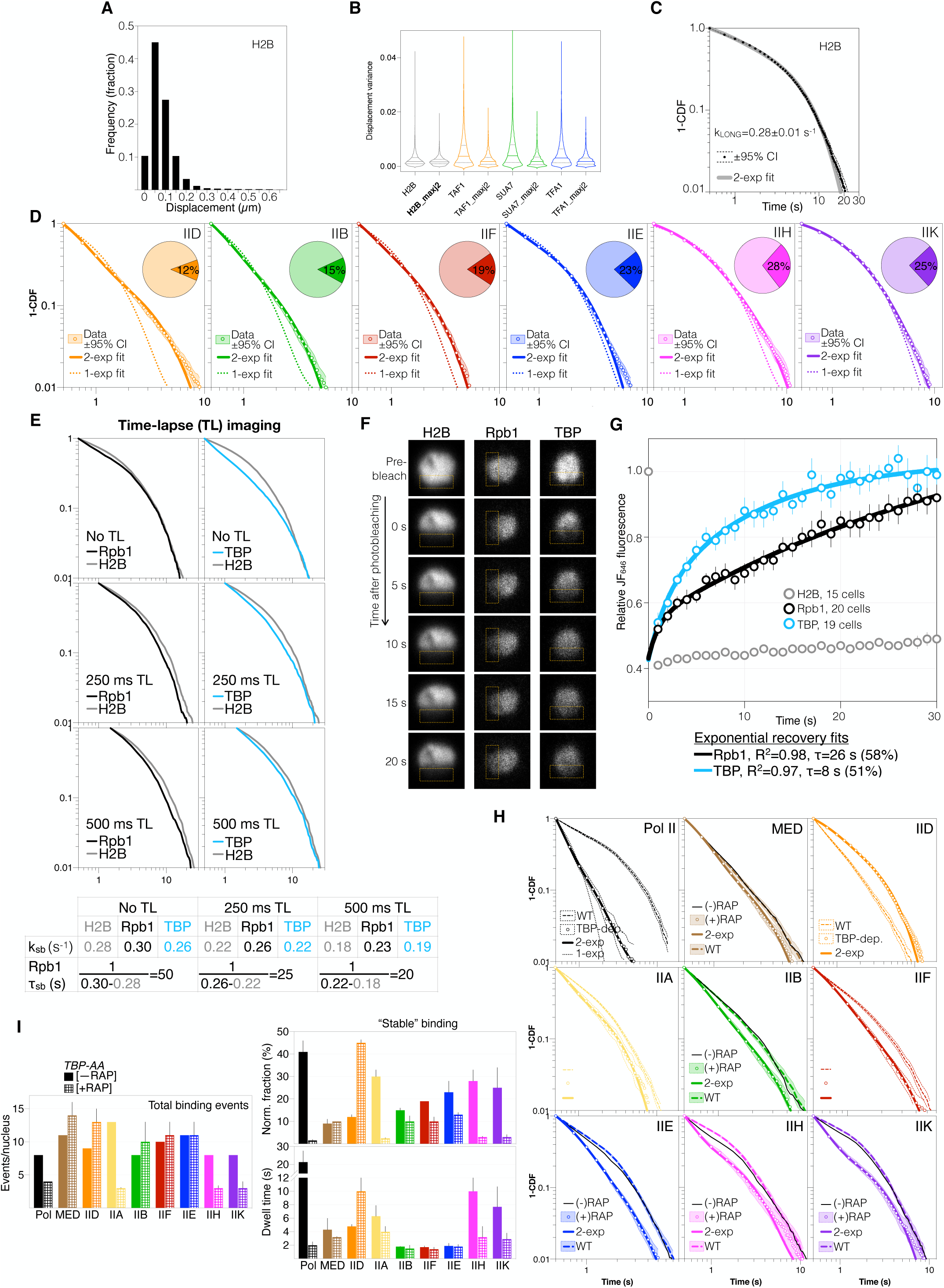
Analyzing slow-tracking and FRAP data, Related to Figure 3. (A) Displacement distribution of H2B molecules (Δt=250 ms) showing the motion of chromatin detected by slow tracking. The histogram indicates the predominance of short displacements rarely exceeding 0.2-0.3 µm (∼2-3 pixels, 1 pixel = 107 nm). (B) Violin plots of variance among displacements by single molecules tracked using a two or three-pixel maximum jump cutoff (see Methods). Trajectories obtained using the two-pixel cutoff exhibit displacement variances similar to that of H2B. (C) Log-log 1-CDF curve for H2B, as in (a). The apparent dissociation constant for stable binding k_sb_ from double-exponential fit, used to correct k_sb_ values for PIC components, is indicated. Curve contains data from three biological replicates. (D) Log-log survival probability (1-CDF) (dots) computed from apparent dwell times of single-molecule binding events. Re-sampling by bootstrapping provided ±95% confidence interval (CI) (dashed lines). Double-exponential fit (solid line) indicated fractions of stable (f_sb_) and transient binding (pie chart, f_sb_ shown) and respective apparent unbinding rates (k_sb_ and k_tb_, Table 1). Curves contain data from three biological replicates. Fit results are reported in Table S1. (E) 1-CDF curves for H2B, Rpb1, and TBP acquired by time-lapse imaging (TL) featuring 250 ms exposure time alternating with 250 ms or 500 ms dark time. Results from double-exponential fitting are shown in bottom table. Similar corrected Rpb1 τ_sb_ were obtained from both TL regimes. However, TBP k_sb_ remained too close that that of H2B for correction. (F) Raw FRAP images of Halo-H2B, Rpb1-Halo, and Halo-TBP labeled with JF^646^, showing nuclear fluorescence before (pre-bleach) and up to 20 s after bleaching. The bleach areas are indicated (rectangles). (G) Quantitation of fluorescence recovery and corresponding fits for Rpb1 and TBP. Fit quality (R^2^) and slow-recovery τ and fraction are indicated for each factor. Data are mean ± s.e.m. from 15 (H2B), 19 (TBP) and 20 cells (Rpb1). The faster TBP recovery is unexpected in light of time-lapse SMT results in (E) and may reflect widely different TBP turnover rates *in vivo*. TBP is highly enriched at Pol I-transcribed genes, where its slow turnover kinetics (Grimaldi et al., 2014; Werven et al., 2009) might be preferentially captured by time-lapse SMT. Further FRAP experiments where the nucleolus, the center of Pol I transcription, is targeted for bleaching may address the discrepancy in these results. (H) Slow-tracking results for *tbp-AA* (-) and (+) RAP conditions. Comparing *tbp-AA* (-)RAP and WT curves indicates similar dissociation kinetics. Double-exponential fits are shown for (+) RAP curves and results reported in Table S2. Curves contain data from one (-RAP) or three (+RAP and WT) biological replicates and are shown with ±95% CI as in (D). (I) Left: Total number of detected binding events (stable and transient) per imaged nucleus under (-) or (+)RAP condition (see Methods). Right: fraction of longer-dwell binding events, normalized for events/nucleus (see Methods) and corrected dwell times.

**Figure S5.**
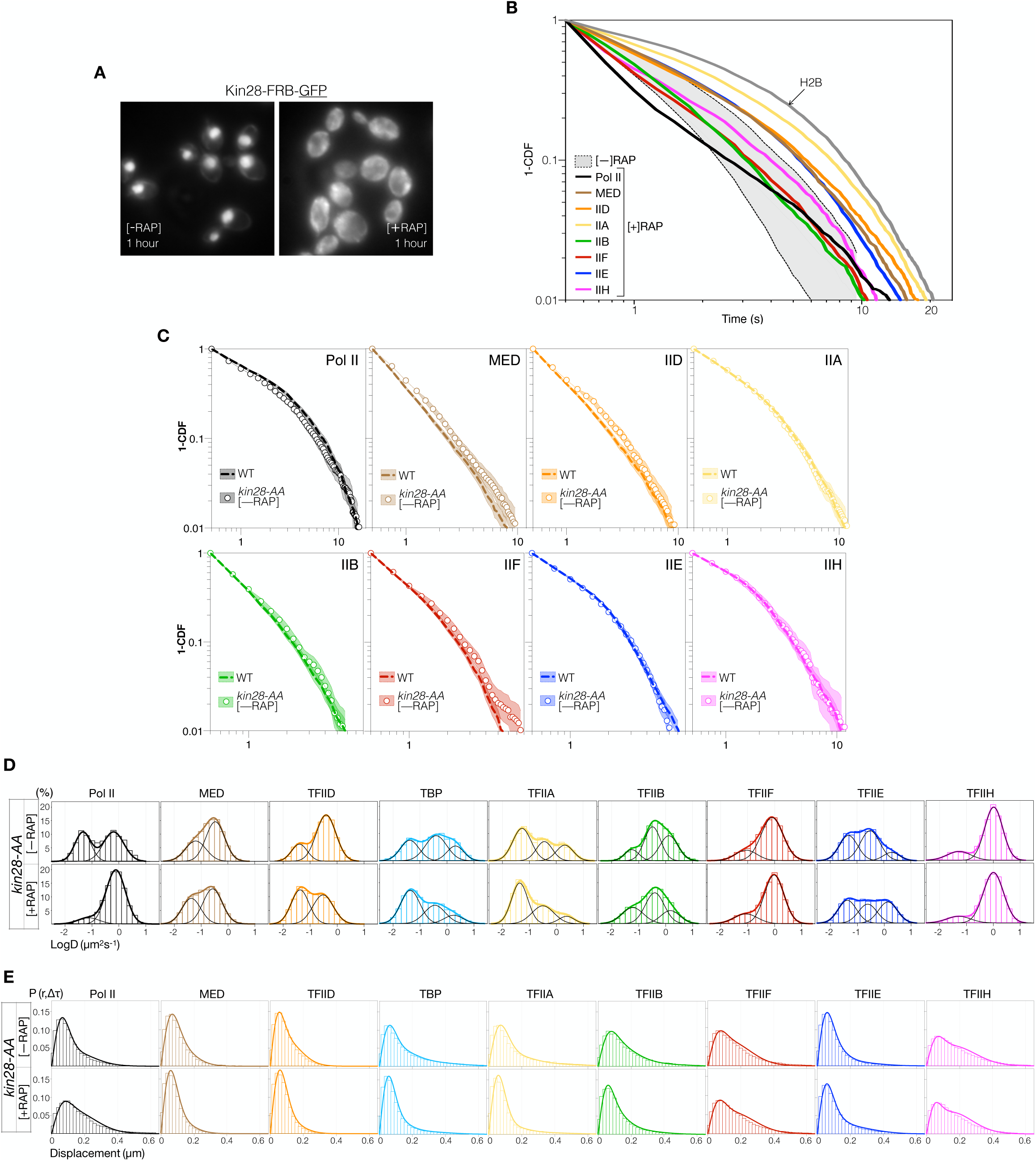
Effect of Kin28 depletion on PIC dynamics, Related to Figure 5. (A) GFP fluorescence images of Kin28 in *kin28-AA* cells without (-RAP, left) and with (+RAP, right) 1 µg/mL rapamycin for 1 hour. (B) Overlay of *kin28-AA* (+RAP) 1-CDF curves from Figure 4C showing extended chromatin residence of Mediator and all GTFs upon Kin28 depletion, compared to the normal range from WT (shaded region). The TFIIA curve approached the H2B limit, precluding reliable photobleaching correction. Thus, we did not quantify and report its dissociation kinetics under this condition. (C) Overlay of *kin28-AA* (-RAP) and WT 1-CDF curves for each PIC component showing similar dissociation kinetics. *kin28-AA* and WT data were obtained from one and three biological replicates, respectively. All curves are shown with ±95% CI. (D) LogD histograms of PIC components in *kin28-AA* cells, shown as in Figure S3C. (E) Kinetic modeling of data in (D). Results are reported in Table S2.

**Figure S6.**
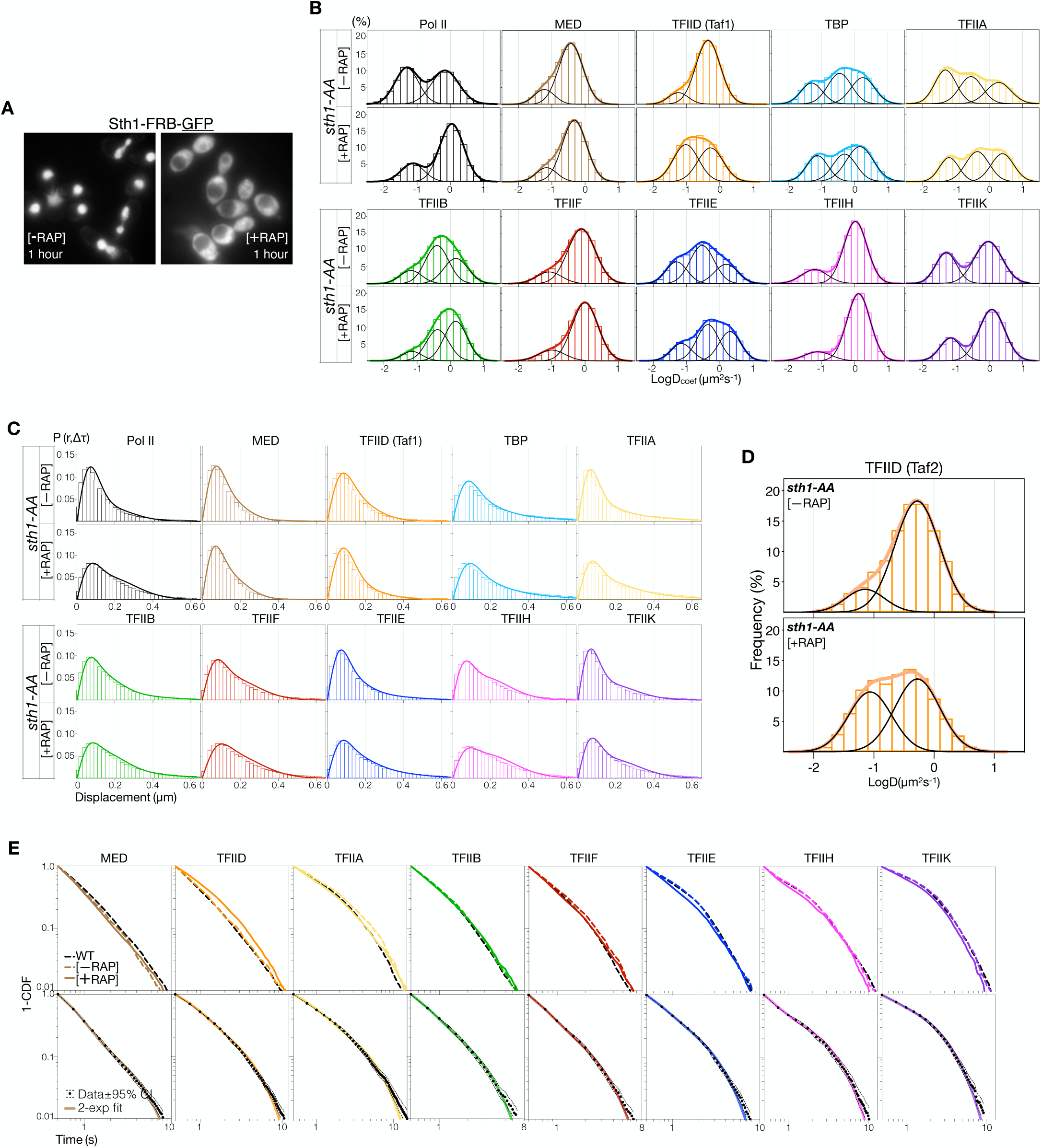
Effect of RSC inactivation on PIC dynamics, Related to Figure 6. (A) GFP fluorescence images of Sth1 in *sth1-AA* cells without (-RAP, left) and with (+RAP, right) 1 µg/mL rapamycin for 1 hour. (B) LogD histograms of PIC components in *sth1-AA* cells, shown as in Figure S3C. All histograms contain data from three biological replicates. (C) Kinetic modeling of data in (B). Results are reported in Table S2. (D) LogD histograms of Taf2 subunit of TFIID showing increased chromatin-bound fraction after RSC inactivation. (E) Top: 1-CDF curves for *sth1-AA* (-) and (+) RAP. Comparing *sth1-AA* (-RAP) and WT curves indicates little effect of the *sth1-AA* genetic background on dissociation kinetics of PIC components. Bottom: *sth1-AA* (+RAP) curves, shown with ±95% CI, and corresponding double-exponential fits. Fit results are reported in Table S2. Data were obtained from one (-RAP) and three (+RAP and WT) biological replicates.

**Figure S7.**
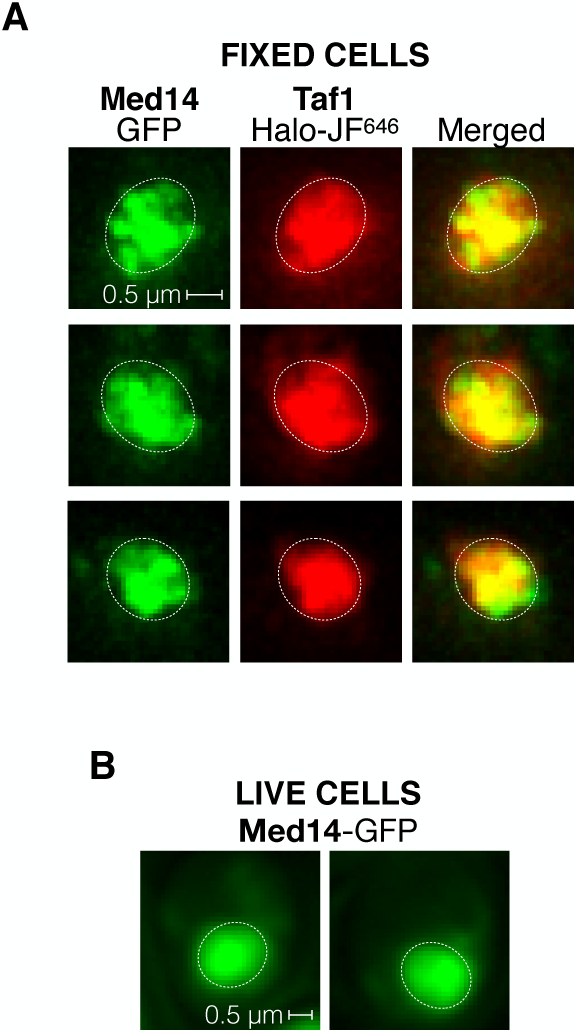
Nuclear distributions of Mediator and TFIID, Related to Figure 7. (A) Two-color deconvolution microscopy images of Med14-GFP (left) and Taf1-Halo-JF^646^ (middle) in three fixed cells showing non-homogenous nuclear distributions. Overlay images (right) show the degree of spatial overlap between prominent Mediator and TFIID foci in the same cells. The approximate nuclear periphery (dashed ovals), based on the overlay image, is indicated for each cell. (B) Med14-GFP images obtained from two live cells showing intense signals in subnuclear regions consistent in dimensions with fixed-cell images in (A).

**Table S1.** Fast- and slow-tracking results for PIC components in wildtype yeast. Related to Figures 1, S1E, 3 and 5. Fit results from Spot-On kinetic modeling of displacements obtained by fast tracking. Average diffusion coefficients (D) and corresponding fractions of molecules (F) are indicated for distinct populations. Trajectories with at least 3 detections (# traj.) were accounted for in the kinetic model (Figure S1E). Data represent mean ± SD from two or three biological replicates (number of *). For slow-tracking data, double-exponential fitting of the 1-CDF survival curves (Figures 3A, 3B and S4D) was performed to obtain the dissociation rate (k) and corresponding fraction (f) of stably-bound (sb) or transiently-bound (tb) molecules. The dwell times τ_sb_ and τ_tb_ resulted from correction using k_sb_ for H2B (see Methods). The total number of binding events (transient and stable) from three biological replicates is indicated for each factor. Fast- and slow-tracking data allowed calculations of search kinetics and occupancy according to Figure 5 and STAR Methods.

**Table S2.** Fast- and slow-tracking results for PIC components in AA strains. Related to Figures 2, 4 and 6. Fast- and slow-tracking, when applicable, results from AA experiments, outlined as in Table S1. For *tbp-AA*, *rpb1-AA* and *kin28-AA*, fast-tracking data were collected for one biological replicate under [-RAP] conditions to verify similar dynamics compared to wildtype. For *tbp-AA*, *kin28-AA* and *sth1-AA*, slow-tracking data were collected for one biological replicate under [-RAP] conditions and corresponding 1-CDF curves shown in Figures S4H, S5C and S6E, respectively. Curves were not fitted.

**Table S3.** S. cervisiae strains used in this study.

**Video S1. Taf1-Halo motion by fast tracking in a live cell, Related to** **Figure 1**. Raw movie of Taf1-Halo-JF^552^ in the nucleus of a live cell. Data was acquired at 10 ms framerate and rendered at 50 fps (frames per second). The nuclear outline (oval) was estimated from total nuclear fluorescence (not shown).

**Video S2. Kin28-Halo motion by slow tracking in a live cell, Related to** **Figure 3**. Raw movie of Kin28-Halo-JF^552^ in the nucleus of a live cell. Two single binding events intervened by a motion-blurred diffusive episode, likely by the same molecule, are captured. Data was acquired at 250 ms framerate and rendered at 25 fps (frames per second). The nuclear outline (oval) was estimated from total nuclear fluorescence (not shown).

## REFERENCES

Ansari, S.A., He, Q., and Morse, R.H. (2009). Mediator complex association with constitutively transcribed genes in yeast. Proc National Acad Sci 106, 16734–16739.

Baek, H.J., Malik, S., Qin, J., and Roeder, R.G. (2002). Requirement of TRAP/Mediator for Both Activator-Independent and Activator-Dependent Transcription in Conjunction with TFIID-Associated TAFIIs. Mol Cell Biol 22, 2842–2852.

Baek, H.J., Kang, Y.K., and Roeder, R.G. (2006). Human Mediator Enhances Basal Transcription by Facilitating Recruitment of Transcription Factor IIB during Preinitiation Complex Assembly. Journal of Biological Chemistry 281, 15172–15181.

Ball, D.A., Mehta, G.D., Salomon-Kent, R., Mazza, D., Morisaki, T., Mueller, F., McNally, J.G., and Karpova, T.S. (2016). Single molecule tracking of Ace1p in Saccharomyces cerevisiae defines a characteristic residence time for non-specific interactions of transcription factors with chromatin. Nucleic Acids Research 44, gkw744.

Baptista, T., Grünberg, S., Minoungou, N., Koster, M.J.E., Timmers, H.T.M., Hahn, S., Devys, D., and Tora, L. (2017). SAGA Is a General Cofactor for RNA Polymerase II Transcription. Molecular Cell 68, 130–143.e5.

Boija, A., Klein, I.A., Sabari, B.R., Dall’Agnese, A., Coffey, E.L., Zamudio, A.V., Li, C.H., Shrinivas, K., Manteiga, J.C., Hannett, N.M., et al. (2018). Transcription Factors Activate Genes through the Phase-Separation Capacity of Their Activation Domains. Cell 175, 1842–1855.e16.

Booher, K.R., and Kaiser, P. (2008). A PCR-based strategy to generate yeast strains expressing endogenous levels of amino-terminal epitope-tagged proteins.

Borggrefe, T., Davis, R., Bareket-Samish, A., and Kornberg, R.D. (2001). Quantitation of the RNA polymerase II transcription machinery in yeast. The Journal of Biological Chemistry 276, 47150–47153.

Brahma, S., and Henikoff, S. (2018). RSC-Associated Subnucleosomes Define MNase-Sensitive Promoters in Yeast. Molecular Cell 73, 238–249.e3.

Chen, J., Zhang, Z., Li, L., Chen, B.-C., Revyakin, A., Hajj, B., Legant, W., Dahan, M., Lionnet, T., Betzig, E., et al. (2014). Single-Molecule Dynamics of Enhanceosome Assembly in Embryonic Stem Cells. Cell 156, 1274–1285.

Cherry, J.M., Adler, C., Ball, C., Chervitz, S.A., Dwight, S.S., Hester, E.T., Jia, Y., Juvik, G., Roe, T., Schroeder, M., et al. (1998). SGD: Saccharomyces Genome Database. Nucleic Acids Res 26, 73–79.

Cho, W.-K., Spille, J.-H., Hecht, M., Lee, C., Li, C., Grube, V., and Cisse, I.I. (2018). Mediator and RNA polymerase II clusters associate in transcription-dependent condensates. Science 361, 412–415.

Cisse, I.I., Izeddin, I., Causse, S.Z., Boudarene, L., Senecal, A., Muresan, L., Dugast-Darzacq, C., Hajj, B., Dahan, M., and Darzacq, X. (2013). Real-Time Dynamics of RNA Polymerase II Clustering in Live Human Cells. Science 341, 664–667.

Compe, E., Genes, C.M., Braun, C., Coin, F., and Egly, J.-M. (2019). TFIIE orchestrates the recruitment of the TFIIH kinase module at promoter before release during transcription. Nature Communications 10, 2084.

Cramer, P. (2019). Organization and regulation of gene transcription. Nature 573, 45–54.

Dion, V., and Gasser, S.M. (2013). Chromatin movement in the maintenance of genome stability. Cell 152, 1355–1364.

Donczew, R., Warfield, L., Pacheco, D., Erijman, A., and Hahn, S. (2020). Two roles for the yeast transcription coactivator SAGA and a set of genes redundantly regulated by TFIID and SAGA. Elife 9, e50109.

Donovan, B.T., Huynh, A., Ball, D.A., Patel, H.P., Poirier, M.G., Larson, D.R., Ferguson, M.L., and Lenstra, T.L. (2019). Live-cell imaging reveals the interplay between transcription factors, nucleosomes, and bursting. The EMBO Journal 38, e51794.

Efron, B., and Tibshirani, R. (1986). Bootstrap Methods for Standard Errors, Confidence Intervals, and Other Measures of Statistical Accuracy. Stat Sci 1, 54–75.

Fujiwara, R., Damodaren, N., Wilusz, J.E., and Murakami, K. (2019). The capping enzyme facilitates promoter escape and assembly of a follow-on preinitiation complex for reinitiation. Proceedings of the National Academy of Sciences 116, 22573–22582.

Ganguli, D., Chereji, R.V., Iben, J.R., Cole, H.A., and Clark, D.J. (2014). RSC-dependent constructive and destructive interference between opposing arrays of phased nucleosomes in yeast. Genome Research 24, 1637–1649.

Gebhardt, J.C.M., Suter, D.M., Roy, R., Zhao, Z.W., Chapman, A.R., Basu, S., Maniatis, T., and Xie, X.S. (2013). Single-molecule imaging of transcription factor binding to DNA in live mammalian cells. Nature Methods 10, 421–426.

Gietz, R.D., and Schiestl, R.H. (2007). High-efficiency yeast transformation using the LiAc/SS carrier DNA/PEG method. Nature Protocols 2, 31–34.

Grimaldi, Y., Ferrari, P., and Strubin, M. (2014). Independent RNA polymerase II preinitiation complex dynamics and nucleosome turnover at promoter sites in vivo. Genome Research 24, 117–124.

Grimm, J.B., English, B.P., Chen, J., Slaughter, J.P., Zhang, Z., Revyakin, A., Patel, R., Macklin, J.J., Normanno, D., Singer, R.H., et al. (2015). A general method to improve fluorophores for live-cell and single-molecule microscopy. Nature Methods 12, 244–50-3 p following 250.

Grünberg, S., Henikoff, S., Hahn, S., and Zentner, G.E. (2016). Mediator binding to UASs is broadly uncoupled from transcription and cooperative with TFIID recruitment to promoters. The EMBO Journal 35, 2435–2446.

Hammond, C.M., Strømme, C.B., Huang, H., Patel, D.J., and Groth, A. (2017). Histone chaperone networks shaping chromatin function. Nature Reviews Molecular Cell Biology 18, 141–158.

Hansen, A.S., Pustova, I., Cattoglio, C., Tjian, R., and Darzacq, X. (2017). CTCF and cohesin regulate chromatin loop stability with distinct dynamics. ELife 6, 2848.

Hansen, A.S., Woringer, M., Grimm, J.B., Lavis, L.D., Tjian, R., and Darzacq, X. (2018). Robust model-based analysis of single-particle tracking experiments with Spot-On. ELife 7.

Hansen, A.S., Amitai, A., Cattoglio, C., Tjian, R., and Darzacq, X. (2020). Guided nuclear exploration increases CTCF target search efficiency. Nat Chem Biol 16, 257–266.

Hartley, P.D., and Madhani, H.D. (2009). Mechanisms that Specify Promoter Nucleosome Location and Identity. Cell 137, 445–458.

Haruki, H., Nishikawa, J., and Laemmli, U.K. (2008). The anchor-away technique: rapid, conditional establishment of yeast mutant phenotypes. Molecular Cell 31, 925–932.

Hirschhorn, J.N., Bortvin, A.L., Ricupero-Hovasse, S.L., and Winston, F. (1995). A new class of histone H2A mutations in Saccharomyces cerevisiae causes specific transcriptional defects in vivo. Mol Cell Biol 15, 1999–2009.

Ho, B., Baryshnikova, A., and Brown, G.W. (2018). Unification of Protein Abundance Datasets Yields a Quantitative Saccharomyces cerevisiae Proteome. Cell Systems 6, 192–205.e3.

Izeddin, I., Récamier, V., Bosanac, L., Cissé, I.I., Boudarene, L., Dugast-Darzacq, C., Proux, F., Bénichou, O., Voituriez, R., Bensaude, O., et al. (2014). Single-molecule tracking in live cells reveals distinct target-search strategies of transcription factors in the nucleus. ELife 3.

Jeronimo, C., Langelier, M.-F., Bataille, A.R., Pascal, J.M., Pugh, B.F., and Robert, F. (2016). Tail and Kinase Modules Differently Regulate Core Mediator Recruitment and Function In Vivo. Molecular Cell 64, 455–466.

Johnson, K.M., and Carey, M. (2003). Assembly of a Mediator/TFIID/TFIIA Complex Bypasses the Need for an Activator. Curr Biol 13, 772–777.

Johnson, K.M., Wang, J., Smallwood, A., Arayata, C., and Carey, M. (2002). TFIID and human mediator coactivator complexes assemble cooperatively on promoter DNA. Genes & Development 16, 1852–1863.

Joo, Y.J., Ficarro, S.B., Soares, L.M., Chun, Y., Marto, J.A., and Buratowski, S. (2017). Downstream promoter interactions of TFIID TAFs facilitate transcription reinitiation. Genes & Development 31, 2162–2174.

Kent, S., Brown, K., Yang, C., Alsaihati, N., Tian, C., Wang, H., and Ren, X. (2020). Phase-Separated Transcriptional Condensates Accelerate Target-Search Process Revealed by Live-Cell Single-Molecule Imaging. Cell Reports 33, 108248.

Keogh, M.-C., Cho, E.-J., Podolny, V., and Buratowski, S. (2002). Kin28 Is Found within TFIIH and a Kin28-Ccl1-Tfb3 Trimer Complex with Differential Sensitivities to T-Loop Phosphorylation. Mol Cell Biol 22, 1288–1297.

Khattabi, L.E., Zhao, H., Kalchschmidt, J., Young, N., Jung, S., Blerkom, P.V., Kieffer-Kwon, P., Kieffer-Kwon, K.-R., Park, S., Wang, X., et al. (2019). A Pliable Mediator Acts as a Functional Rather Than an Architectural Bridge between Promoters and Enhancers. Cell 178, 1145–1158.e20.

Kimura, H., Tao, Y., Roeder, R.G., and Cook, P.R. (1999). Quantitation of RNA Polymerase II and Its Transcription Factors in an HeLa Cell: Little Soluble Holoenzyme but Significant Amounts of Polymerases Attached to the Nuclear Substructure. Molecular and Cellular Biology 19, 5383–5392.

Klein-Brill, A., Joseph-Strauss, D., Appleboim, A., and Friedman, N. (2019). Dynamics of Chromatin and Transcription during Transient Depletion of the RSC Chromatin Remodeling Complex. Cell Reports 26, 279–292.e5.

Knoll, E.R., Zhu, Z.I., Sarkar, D., Landsman, D., and Morse, R.H. (2018). Role of the pre-initiation complex in Mediator recruitment and dynamics. ELife 7.

Knoll, E.R., Zhu, Z.I., Sarkar, D., Landsman, D., and Morse, R.H. (2020). Kin28 depletionincreases association of TFIID subunits Taf1 and Taf4 with promoters in Saccharomyces cerevisiae. Nucleic Acids Res 48, 4244–4255.

Kraemer, S.M., Ranallo, R.T., Ogg, R.C., and Stargell, L.A. (2001). TFIIA Interacts with TFIID via Association with TATA-Binding Protein and TAF40. Mol Cell Biol 21, 1737– 1746.

Kubik, S., Bruzzone, M.J., Jacquet, P., Falcone, J.-L., Rougemont, J., and Shore, D. (2015). Nucleosome Stability Distinguishes Two Different Promoter Types at All Protein-Coding Genes in Yeast. Molecular Cell 60, 422–434.

Kubik, S., O’Duibhir, E., Jonge, W.J. de, Mattarocci, S., Albert, B., Falcone, J.-L., Bruzzone, M.J., Holstege, F.C.P., and Shore, D. (2018). Sequence-Directed Action of RSC Remodeler and General Regulatory Factors Modulates +1 Nucleosome Position to Facilitate Transcription. Molecular Cell 71, 89–102.e5.

Lauberth, S.M., Nakayama, T., Wu, X., Ferris, A.L., Tang, Z., Hughes, S.H., and Roeder, R.G. (2013). H3K4me3 Interactions with TAF3 Regulate Preinitiation Complex Assembly and Selective Gene Activation. Cell 152, 1021–1036.

Layer, J.H., and Weil, P.A. (2013). Direct TFIIA-TFIID protein contacts drive budding yeast ribosomal protein gene transcription. The Journal of Biological Chemistry 288, 23273– 23294.

Le, S.N., Brown, C.R., Harvey, S., Boeger, H., Elmlund, H., and Elmlund, D. (2019). The TAFs of TFIID Bind and Rearrange the Topology of the TATA-Less RPS5 Promoter. International Journal of Molecular Sciences 20, 3290.

Lerner, J., Gomez-Garcia, P.A., McCarthy, R.L., Liu, Z., Lakadamyali, M., and Zaret, K.S. (2020). Two-Parameter Mobility Assessments Discriminate Diverse Regulatory Factor Behaviors in Chromatin. Mol Cell.

Li, J., Dong, A., Saydaminova, K., Chang, H., Wang, G., Ochiai, H., Yamamoto, T., and Pertsinidis, A. (2019). Single-Molecule Nanoscopy Elucidates RNA Polymerase II Transcription at Single Genes in Live Cells. Cell 178, 491–506.e28.

Lim, M.K., Tang, V., Saux, A.L., Schüller, J., Bongards, C., and Lehming, N. (2007). Gal11p Dosage-compensates Transcriptional Activator Deletions via Taf14p. J Mol Biol 374, 9–23.

Loffreda, A., Jacchetti, E., Antunes, S., Rainone, P., Daniele, T., Morisaki, T., Bianchi, M.E., Tacchetti, C., and Mazza, D. (2017). Live-cell p53 single-molecule binding is modulated by C-terminal acetylation and correlates with transcriptional activity. Nature Communications 8, 313.

Maxon, M.E., Goodrich, J.A., and Tjian, R. (1994). Transcription factor IIE binds preferentially to RNA polymerase IIa and recruits TFIIH: a model for promoter clearance. Gene Dev 8, 515–524.

Mazza, D., Abernathy, A., Golob, N., Morisaki, T., and McNally, J.G. (2012). A benchmark for chromatin binding measurements in live cells. Nucleic Acids Research 40, e119.

McSwiggen, D.T., Hansen, A.S., Teves, S.S., Marie-Nelly, H., Hao, Y., Heckert, A.B., Umemoto, K.K., Dugast-Darzacq, C., Tjian, R., and Darzacq, X. (2019). Evidence for DNA-mediated nuclear compartmentalization distinct from phase separation. Elife 8, e47098.

Mehta, G.D., Ball, D.A., Eriksson, P.R., Chereji, R.V., Clark, D.J., McNally, J.G., and Karpova, T.S. (2018). Single-Molecule Analysis Reveals Linked Cycles of RSC Chromatin Remodeling and Ace1p Transcription Factor Binding in Yeast. Molecular Cell 72, 875–887.e9.

Miné-Hattab, J., Recamier, V., Izeddin, I., Rothstein, R., and Darzacq, X. (2017). Multi-scale tracking reveals scale-dependent chromatin dynamics after DNA damage. Mol Biol Cell 28, 3323–3332.

Murakami, K., Mattei, P.-J., Davis, R.E., Jin, H., Kaplan, C.D., and Kornberg, R.D. (2015). Uncoupling Promoter Opening from Start-Site Scanning. Molecular Cell 59, 133–138.

Neil, H., Malabat, C., d’Aubenton-Carafa, Y., Xu, Z., Steinmetz, L.M., and Jacquier, A. (2009). Widespread bidirectional promoters are the major source of cryptic transcripts in yeast. Nature 457, 1038–1042.

Normanno, D., Dahan, M., and Darzacq, X. (2012). Intra-nuclear mobility and target search mechanisms of transcription factors: A single-molecule perspective on gene expression. Biochimica et Biophysica Acta (BBA) - Gene Regulatory Mechanisms 1819, 482–493.

Papai, G., Tripathi, M.K., Ruhlmann, C., Layer, J.H., Weil, P.A., and Schultz, P. (2010). TFIIA and the transactivator Rap1 cooperate to commit TFIID for transcription initiation. Nature 465, 956–960.

Patel, A.B., Louder, R.K., Greber, B.J., Grünberg, S., Luo, J., Fang, J., Liu, Y., Ranish, J., Hahn, S., and Nogales, E. (2018). Structure of human TFIID and mechanism of TBP loading onto promoter DNA. Science 362, eaau8872.

Pelechano, V., Chávez, S., and Pérez-Ortín, J.E. (2010). A Complete Set of Nascent Transcription Rates for Yeast Genes. PLoS ONE 5, e15442.

Petrenko, N., Jin, Y., Wong, K.H., and Struhl, K. (2016). Mediator Undergoes a Compositional Change during Transcriptional Activation. Molecular Cell 64, 443–454.

Petrenko, N., Jin, Y., Wong, K.H., and Struhl, K. (2017). Evidence that Mediator is essential for Pol II transcription, but is not a required component of the preinitiation complex in vivo. ELife 6.

Qiu, C., Jin, H., Vvedenskaya, I., Llenas, J.A., Zhao, T., Malik, I., Schwartz, S.L., Cui, P., Čabart, P., Han, K.H., et al. (2019). Promoter scanning during transcription initiation in Saccharomyces cerevisiae: Pol II in the “shooting gallery”. Biorxiv 810127.

Ramachandran, S., Zentner, G.E., and Henikoff, S. (2015). Asymmetric nucleosomes flank promoters in the budding yeast genome. Genome Research 25, 381–390.

Rani, P.G., Ranish, J.A., and Hahn, S. (2004). RNA Polymerase II (Pol II)-TFIIF and Pol II-Mediator Complexes: the Major Stable Pol II Complexes and Their Activity in Transcription Initiation and Reinitiation. Mol Cell Biol 24, 1709–1720.

Rhee, H.S., and Pugh, B.F. (2012). Genome-wide structure and organization of eukaryotic pre-initiation complexes. Nature 483, 295–301.

Robinson, P.J., Trnka, M.J., Bushnell, D.A., Davis, R.E., Mattei, P.-J., Burlingame, A.L., and Kornberg, R.D. (2016). Structure of a Complete Mediator-RNA Polymerase II Pre-Initiation Complex. Cell 166, 1411–1422.e16.

Rodríguez-Molina, J.B., Tseng, S.C., Simonett, S.P., Taunton, J., and Ansari, A.Z. (2016). Engineered Covalent Inactivation of TFIIH-Kinase Reveals an Elongation Checkpoint and Results in Widespread mRNA Stabilization. Molecular Cell 63, 433–444.

Schilbach, S., Hantsche, M., Tegunov, D., Dienemann, C., Wigge, C., Urlaub, H., and Cramer, P. (2017). Structures of transcription pre-initiation complex with TFIIH and Mediator. Nature 551, 204–209.

Schindelin, J., Arganda-Carreras, I., Frise, E., Kaynig, V., Longair, M., Pietzsch, T., Preibisch, S., Rueden, C., Saalfeld, S., Schmid, B., et al. (2012). Fiji: an open-source platform for biological-image analysis. Nat Methods 9, 676–682.

Schweikhard, V., Meng, C., Murakami, K., Kaplan, C.D., Kornberg, R.D., and Block, S.M. (2014). Transcription factors TFIIF and TFIIS promote transcript elongation by RNA polymerase II by synergistic and independent mechanisms. Proc National Acad Sci 111, 6642–6647.

Shrinivas, K., Sabari, B.R., Coffey, E.L., Klein, I.A., Boija, A., Zamudio, A.V., Schuijers, J., Hannett, N.M., Sharp, P.A., Young, R.A., et al. (2019). Enhancer Features that Drive Formation of Transcriptional Condensates. Molecular Cell 75, 549–561.e7.

Sprouse, R.O., Karpova, T.S., Mueller, F., Dasgupta, A., McNally, J.G., and Auble, D.T. (2008). Regulation of TATA-binding protein dynamics in living yeast cells. PNAS 105, 13304–13308.

Tatavosian, R., Duc, H.N., Huynh, T.N., Fang, D., Schmitt, B., Shi, X., Deng, Y., Phiel, C., Yao, T., Zhang, Z., et al. (2018). Live-cell single-molecule dynamics of PcG proteins imposed by the DIPG H3.3K27M mutation. Nature Communications 9, 2080.

Thevenaz, P., Ruttimann, U.E., and Unser, M. (1998). A pyramid approach to subpixel registration based on intensity. Ieee T Image Process 7, 27–41.

Tuttle, L.M., Pacheco, D., Warfield, L., Luo, J., Ranish, J., Hahn, S., and Klevit, R.E. (2018). Gcn4-Mediator Specificity Is Mediated by a Large and Dynamic Fuzzy Protein-Protein Complex. Cell Reports 22, 3251–3264.

Vallotton, P., and Olivier, S. (2013). Tri-track: Free Software for Large-Scale Particle Tracking. Microscopy and Microanalysis 19, 451–460.

Vermeulen, M., Mulder, K.W., Denissov, S., Pijnappel, W.W.M.P., Schaik, F.M.A. van, Varier, R.A., Baltissen, M.P.A., Stunnenberg, H.G., Mann, M., and Timmers, H.Th.M. (2007). Selective Anchoring of TFIID to Nucleosomes by Trimethylation of Histone H3 Lysine 4. Cell 131, 58–69.

Warfield, L., Ramachandran, S., Baptista, T., Devys, D., Tora, L., and Hahn, S. (2017). Transcription of Nearly All Yeast RNA Polymerase II-Transcribed Genes Is Dependent on Transcription Factor TFIID. Molecular Cell 68.

Werven, F.J. van, Teeffelen, H.A.A.M. van, Holstege, F.C.P., and Timmers, H.T.M. (2009). Distinct promoter dynamics of the basal transcription factor TBP across the yeast genome. Nature Structural & Molecular Biology 16, 1043–1048.

Wieser, S., and Schütz, G.J. (2008). Tracking single molecules in the live cell plasma membrane—Do’s and Don’t’s. Methods 46, 131–140.

Wong, K.H., Jin, Y., and Struhl, K. (2014). TFIIH phosphorylation of the Pol II CTD stimulates mediator dissociation from the preinitiation complex and promoter escape. Molecular Cell 54, 601–612.

Woringer, M., and Darzacq, X. (2018). Protein motion in the nucleus: from anomalous diffusion to weak interactions. Biochem Soc T 46, BST20170310.

Woringer, M., Darzacq, X., and Izeddin, I. (2014). Geometry of the nucleus: a perspective on gene expression regulation. Current Opinion in Chemical Biology 20, 112–119.

Woringer, M., Izeddin, I., Favard, C., and Berry, H. (2020). Anomalous Subdiffusion in Living Cells: Bridging the Gap Between Experiments and Realistic Models Through Collaborative Challenges. Aip Conf Proc 8, 134.

Xu, Z., Wei, W., Gagneur, J., Perocchi, F., Clauder-Münster, S., Camblong, J., Guffanti, E., Stutz, F., Huber, W., and Steinmetz, L.M. (2009). Bidirectional promoters generate pervasive transcription in yeast. Nature 457, 1033–1037.

Xue, Y., Pradhan, S.K., Sun, F., Chronis, C., Tran, N., Su, T., Van, C., Vashisht, A., Wohlschlegel, J., Peterson, C.L., et al. (2017). Mot1, Ino80C, and NC2 Function Coordinately to Regulate Pervasive Transcription in Yeast and Mammals. Molecular Cell 67, 594–607.e4.

Zawel, L., Kumar, K.P., and Reinberg, D. (1995). Recycling of the general transcription factors during RNA polymerase II transcription. Gene Dev 9, 1479–1490.

Zhang, Z., English, B.P., Grimm, J.B., Kazane, S.A., Hu, W., Tsai, A., Inouye, C., You, C., Piehler, J., Schultz, P.G., et al. (2016). Rapid dynamics of general transcription factor TFIIB binding during preinitiation complex assembly revealed by single-molecule analysis. Genes & Development 30, 2106–2118.

Zheng, Q., Ayala, A.X., Chung, I., Weigel, A.V., Ranjan, A., Falco, N., Grimm, J.B., Tkachuk, A.N., Wu, C., Lippincott-Schwartz, J., et al. (2019). Rational Design of Fluorogenic and Spontaneously Blinking Labels for Super-Resolution Imaging. Acs Central Sci 5, 1602–1613.

